# CLCd and CLCf act redundantly at the TGN/EE and prevent acidification of the Golgi stack

**DOI:** 10.1101/2021.04.07.438841

**Authors:** Stefan Scholl, Stefan Hilmer, Melanie Krebs, Karin Schumacher

## Abstract

The trans-Golgi network/early endosome (TGN/EE) serves as the central hub in which exo- and endocytic trafficking pathways converge and specificity of cargo routing needs to be achieved. Acidification is a hallmark of the TGN/EE and is maintained by the vacuolar H^+^-ATPase (V-ATPase) with support of proton-coupled antiporters. We show here that CLCd and CLCf, two distantly related members of the Arabidopsis chloride channel (CLC)-family that colocalize in the TGN/EE act redundantly and are essential for male gametophyte development. Combining an inducible knock-down approach and in vivo pH-measurements, we show here that reduced CLC-activity does not affect pH in the TGN/EE but causes accumulation of the V-ATPase in trans-Golgi cisternae leading to their hyper-acidification. Taken together, our results show that CLC-mediated anion transport into the TGN/EE is essential and affects spatio-temporal aspects of TGN/EE-maturation as well as its functional separation from the Golgi stack.

## Introduction

Transport through the endomembrane system involves a multitude of interactions between cargo and components of the trafficking machinery. When and where in the endomembrane system reactions including receptor-ligand dissociation, protein processing and protein modification occur is to a large extent controlled by their pH-optimum. Progressive acidification of the secretory and endocytic routes is mediated by the vacuolar-type H^+^-ATPase (V-ATPase), a nano-scale motor that couples the hydrolysis of ATP to the transmembrane movement of protons. Because this activity is electrogenic and generates a transmembrane voltage, for net pumping to occur another ion must move. Cations moving out or anions moving in with protons can serve as counterions that dissipate the voltage. Differential localization of the V-ATPase is mediated by the largest subunit of the membrane-integral Vo-subcomplex (subunit a, ATP6V0a or VHA-a). Isoforms of subunit a are found in all most all eukaryotes and have been shown to target the V-ATPase to the late Golgi (Stv1) and the vacuole (Vph1) in yeast (Kane et al., 2020) and to neurotransmitter vesicles or endosomes (a1), the Golgi (a2), lysosomes (a3) and the plasma membrane (a3, a4) in mammalian cells (Vasanthakumar and Rubinstein, 2020). Within a given compartment the V-ATPase is often found in the presence of a specific member of the chloride channel (CLC)-family of anion transporters. Gef1, the sole yeast CLC-protein is found in the Golgi along with Stv1 (Schwappach et al., 1998) and mammalian CLCs are found in neurotransmitter vesicles (CLC-3), endosomes (CLC-5) and lysosomes (CLC-7; (Jentsch and Pusch, 2018). Given that mutations in either CLCs or V-ATPase subunits lead to similar phenotypes or diseases it was proposed that CLC-proteins might serve as the shunt conductances that enable V-ATPase mediated acidification (Mindell, 2012). Even after the discovery that many members of the CLC-family are secondary active transporters rather than channels (Accardi and Miller, 2004) this hypothesis still seemed plausible as their 2Cl^-^/H^+^ exchange mode would be compatible with a chloride-dependent shunt. However, loss of CLCs does not always impact acidification of the respective compartment (Kasper et al., 2005) and the use of uncoupling mutations that convert the exchangers into a chloride-permeable pore has clearly demonstrated that the capacity for luminal chloride accumulation is a critical function of CLC-proteins (Novarino et al., 2010; Weinert et al., 2020). In plants, chloride is an essential micronutrient required in the oxygen evolving complex of PSII and its toxicity under salt stress conditions has received considerable attention (Raven, 2017). However, even species that are sensitive to high chloride concentrations accumulate much more of it than required for efficient photosynthesis and chloride can thus also be regarded as a beneficial macronutrient that stimulates growth and biomass accumulation due to its osmotic function (Wege et al., 2017). Due to their physico-chemical similarity, the fate of chloride and nitrate, the other abundant inorganic anion found in plant cells, is intimately linked. Understanding the specificity of the transporters involved in uptake, storage and translocation of nitrate and chloride will be crucial in determining the physiological role of chloride more precisely. For members of the plant CLC family, a single amino acid inside the pore region defines their transport selectivity (Wege et al., 2010; Zifarelli and Pusch, 2010). Among the seven CLC-proteins of Arabidopsis, CLCa and CLCb have been shown to be selective for nitrate and mediate its uptake into the vacuole (De Angeli et al., 2006)(von der Fecht-Bartenbach et al., 2010). In addition, it has recently been shown that in guard cells CLCa accounts for cytosolic acidification in response to NO ^-^ (Demes et al., 2020) demonstrating that in addition to mediating the accumulation of anions CLCs that operate as H^+^-coupled exchangers can also contribute to cytosolic pH-homeostasis. According to its selectivity motif CLCc is a tonoplast Cl^-^ transporter predominantly expressed in guard cells and required for turgor changes during stomatal opening (Jossier et al., 2010). The putative chloride channel CLCg also resides at the tonoplast and has been implicated in salt tolerance (Nguyen et al., 2016), whereas CLCe is found in the thylakoid membrane and has been proposed to be involved in nitrite uptake and Cl^-^ homeostasis (Marmagne et al., 2007; Monachello et al., 2009). CLCd has been shown to reside in the trans-Golgi network/early endosome (TGN/EE) where it colocalizes with V-ATPase complexes containing the subunit a isoform VHA-a1 (Dettmer et al., 2006; von der Fecht-Bartenbach et al., 2007). The TGN/EE is the central hub for protein sorting and inhibition of the V-ATPase interferes with endocytic and secretory trafficking (Dettmer et al., 2006; Luo et al., 2015; Viotti et al., 2010). The fact that mutants lacking CLCd display hypersensitivity to the V-ATPase inhibitor Concanamycin A (ConcA) indicates that CLC-activity might be required for acidification of the TGN/EE (Dettmer et al., 2006; von der Fecht-Bartenbach et al., 2007), however a more direct involvement of chloride in the secretory pathway should not be neglected. In both scenarios, the absence of a severe phenotype argues that CLCd might be acting redundantly. Here, we show that CLCf, the only member of the Arabidopsis CLC-family so far largely uncharacterized, colocalizes with VHA-a1 and CLCd in the TGN/EE. The combined loss of CLCd and CLCf causes male gametophyte lethality indicating that CLC-activity in the TGN/EE is essential. By using an artificial miRNA (amiRNA) -based inducible knock-down of CLCf in the *clcd* null background, we show that reduced CLC-activity limits cell expansion. In vivo pH-measurements reveal that pH in the TGN/EE is unaffected by CLC-limitation, however pH in the Golgi stack is substantially lowered.

## Results

### CLCe and CLCf belong to a CLC subgroup conserved in streptophytes

Phylogenetic analysis of CLC protein sequences of the green lineage (*Viridiplantae*) revealed that orthologues of Arabidopsis CLCe and CLCf are present in many species including the charophytes *Chara braunii* and *Klebsormidium nitens* but are absent in the chlorophytes (Subgroup II, Figure 1A). In contrast, orthologues of CLCd are also present in the chlorophytes *Chlamydomonas reinhardtii* and *Chloropicon primus*, whereas a third subclade of the CLC family (CLCv) has representatives from all groups of the viridiplantae but is absent in vascular plants (Subgroups I + III, Figure 1A). Based on its atypical sequence in the region of the selectivity filter and the proposed absence of the so-called proton glutamate, it has been suggested that CLCf could act as a channel (Zifarelli and Pusch, 2010). However, 3D homology modelling using the Cryo-EM structure of chicken CLC-7 (Schrecker et al., 2020) predicts that E247 serves as the gating glutamate, E300 is found in the position of the proton glutamate and that Y463 could be involved in chloride binding (Figure 1B). The fact that all three residues are found among all members of subclade II and are embedded in regions of high sequence conservation (Figure 1C, 1D) supports our conclusion that CLCe and CLCf function are likely to act as antiporters, however in the absence of structural and functional data for members of subgroup II CLCs their anion selectivity remains to be determined.

**Figure 1:**
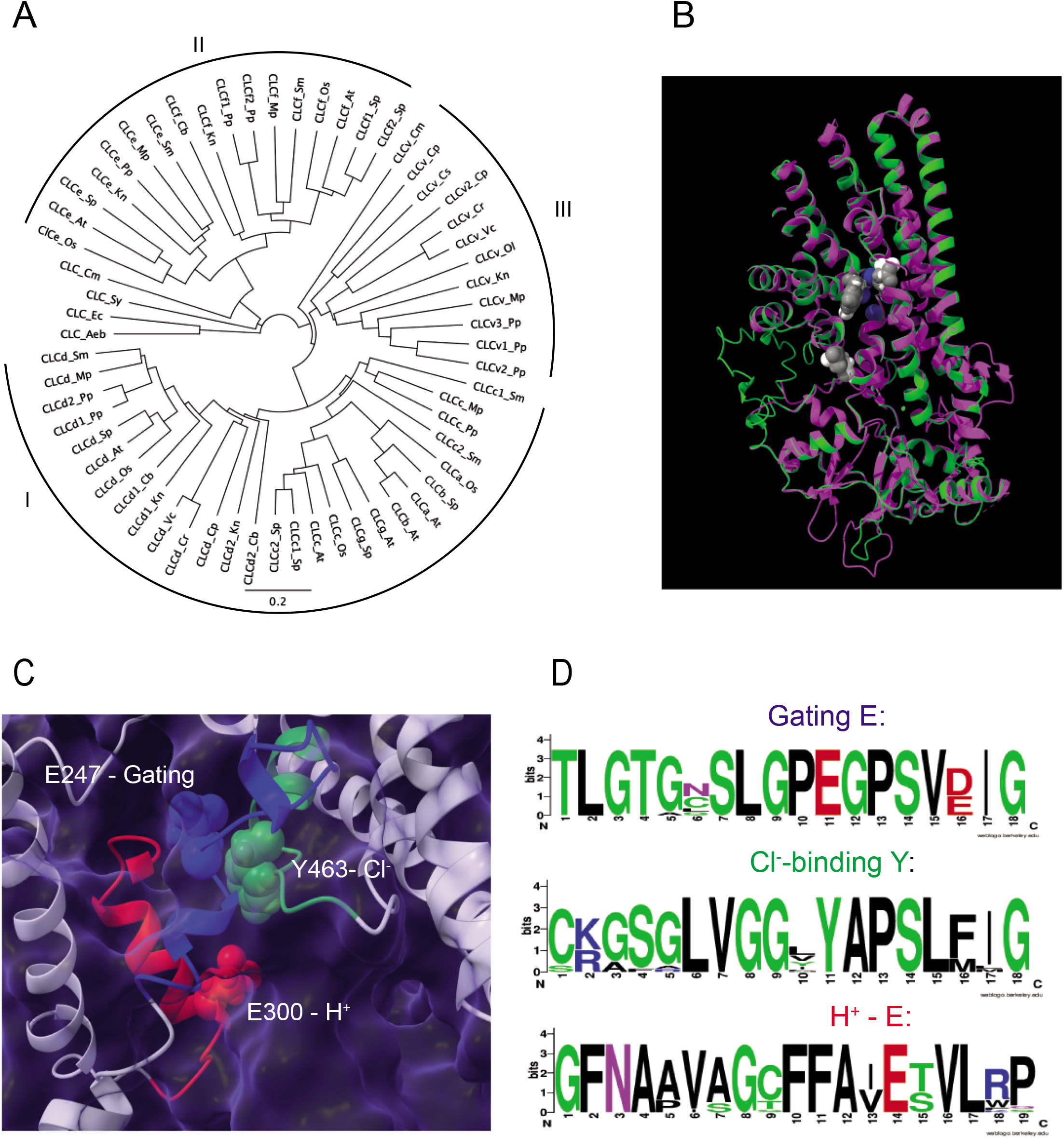
CLCf belongs to CLC-subgroup II and is predicted to be a H^+^/Cl^-^-antiporter. **(A)** Phylogenetic analysis of CLC protein sequences from prokaryotes, a red algae and across the green lineage. Aeb: *Aeromonas bestiarium*, At: *Arabidopsis thaliana*, Cb: *Chara braunii*, Cl: *Chlamydomonas reinhardtii*, Cp: *Chloropicon primus*, Cs: *Coccomyxa subellipsoidea*, Cm: *Cyanidioschyzon merolae*, Ec: *Escherichia coli*, Kn: *Klebsormidium nitens*, Mp: *Marchantia polymorpha*, Os: *Oryza sativa*, Ol: *Ostreococcus lucimarinus*, Pp: *Physcomitrium patens*, Sm: *Selaginella moellendorffii*, Sp: *Solanum pennellii*, Sy: *Synechococus sp*., Vc: *Volvox carteri* **(B)** 3D structure of CLCf based on homology modelling with chicken CLC-7 (7JM6), CLCf is shown in green, CLC-7 in magenta, CLCf E247, E300 and Y463 (grey spheres) are predicted to occupy identical positions a E245, E312 and Y512 (white spheres) in CLC-7. **(C)** Close-up showing the positions of CLCf E247, E300 and Y463 **(D)** Consensus sequences for the corresponding regions across subgroup II CLCs.

### CLCf colocalizes with VHA-a1 and CLCd at the TGN/EE

Based on transient expression in onion epidermal cells CLCf has been reported to be localized at Golgi stacks (Marmagne et al., 2007). To determine the endogenous localisation of CLCf, we fused the full length *CLCf* coding sequence to *mRFP* and expressed the resulting CLCf-mRFP fusion protein under the control of the constitutive *UBQ10* promoter (Grefen et al., 2010). Confocal scanning laser microscopy (CLSM) of stable transgenics coexpressing CLCf-mRFP with the trans-Golgi marker ST-GFP revealed a motile punctate signal pattern for CLCf-mRFP with limited colocalization (Figure 2A). In contrast, when CLCf-mRFP was coexpressed with TGN/EE marker VHA-a1-GFP (Figure 2B) or CLCd-GFP (Figure 2C) the patterns strongly overlapped and quantitative analysis revealed a high degree of colocalization between CLCf-mRFP and VHA-a1-GFP (Figure 2D) comparable to values obtained when VHA-a1-GFP and VHA-a1-mRFP are coexpressed (Figure 2E). Furthermore, gold particles were observed at the TGN/EE when high pressure frozen and freeze substituted seedlings expressing CLCf-mRFP were subjected to immunogold labelling (Figure 2F). As CLCd and CLCf colocalize in the TGN/EE and CLCs are known to form homo-as well as heterodimers (Jentsch and Pusch, 2018; Weinert et al., 2020), we performed Förster resonance energy transfer fluorescence lifetime imaging microscopy (FRET-FLIM) measurements in seedlings coexpressing CLCf-mRFP as FRET acceptor together with CLCd-GFP or VHA-a1-GFP as FRET donors. pHusion-SYP61, in which GFP and mRFP are directly linked (Luo et al., 2015) was used as a FRET positive control. We did not observe differences in GFP lifetime for CLCd-GFP (Supplemental Figure 1A) or VHA-a1-GFP (Supplemental Figure 1B) in the presence or absence of CLCf-mRFP. In conclusion, evidence for heterodimerization between CLCd and CLCf or for a direct interaction between CLCf and VHA-a1 could not be obtained using FRET-FLIM.

**Figure 2:**
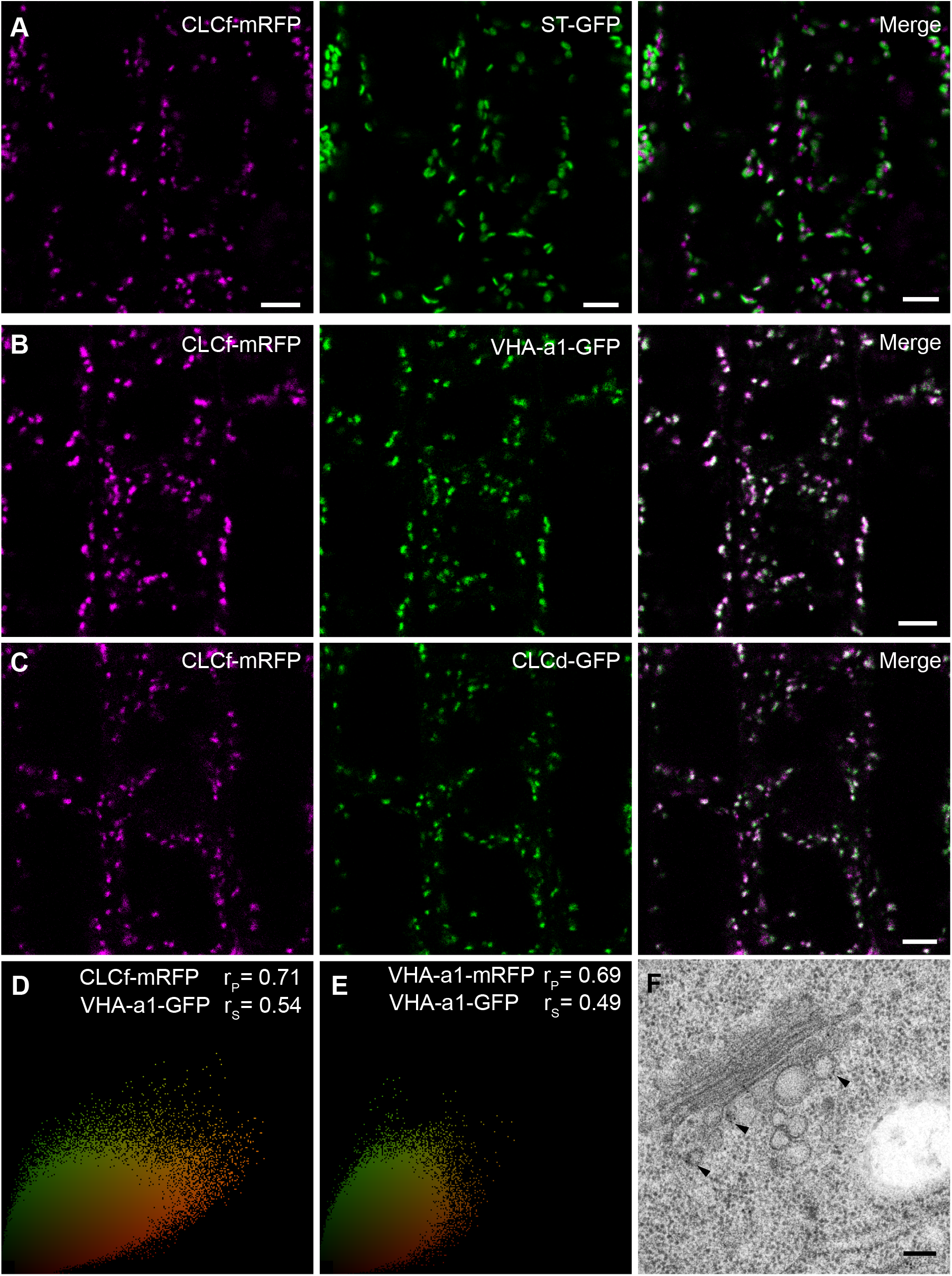
CLCf colocalizes with CLCd and VHA-a1 at the TGN/EE. Localisation analyses was performed in cells of the root elongation zone of 6-day-old Arabidopsis seedlings. **(A)** CLCf-mRFP shows limited overlap with the *trans*-Golgi marker ST-GFP. **(B)** CLCf-mRFP colocalises with the TGN/EE marker VHA-a1-GFP. **(C)** CLCf-mRFP colocalises with CLCd-GFP **(D, E)** Quantification of colocalization for CLCf-mRFP and VHA-a1-GFP **(D)** and for VHA-a1-mRFP and VHA-a1-GFP **(E)** using Pearson (r_p_) and Spearman (r_s_) correlation coefficients. **(F)** Immunogold labelling of high-pressure frozen, freeze substituted Arabidopsis root cells expressing CLCf-mRFP. Sections were labelled with α-DsRed primary antibody and a secondary antibody linked to 10 nm colloidal gold. Arrowheads point to gold particles. Scale bars represents 5 µm in **(A-E)** and 100 nm in **(F)**.

### The presence of CLCs at the TGN/EE is essential

To determine the in vivo function of CLCf, we identified a homozygous T-DNA insertion line from the SALK collection (SALK_112962). PCR-based genotyping and sequencing confirmed a T-DNA insertion in intron 6 of CLCf and PCR on cDNA using primers flanking the insertion site revealed that the T-DNA was not spliced out and full-length transcript could not be detected (Figure 3A). However, similar to *clcd* knock-outs, *clcf* mutant plants are indistinguishable from Col-0 wildtype under standard growth conditions (Figure 3B). To test if *clcf*, like *clcd* (von der Fecht-Bartenbach et al., 2007) displays hypersensitivity to ConcA, we measured hypocotyl length of etiolated seedlings in presence of the V-ATPase inhibitor Concanamycin A (ConcA). Whereas on control medium hypocotyl length in all genotypes was comparable, *clcf* and *clcd* were found to be 15 % shorter than wildtype when grown on 100 µM ConcA (Figure 3C). Lines expressing CLCf-mRFP in the *clcf* background behaved like wildtype indicating that CLCf-mRFP is functional (Figure 3C). Root growth defects on alkaline medium (von der Fecht-Bartenbach et al., 2007) could not be confirmed (Supplementary Figure S2A). We next attempted to identify the *clcd clcf* double mutant by crossing both T-DNA insertion lines. In mature siliques of *clcd/+ clcf* and *clcf clcd/+* plants, roughly 25 % aborted seeds or degenerated ovules were present suggesting that a knock-out of both genes results in lethality (Figure 3D). We performed reciprocal crosses of *clcd clcf/+* with Col-0 to test whether gametophyte development was affected. Via genotyping of F1 offspring of *clcd clcf/+* crossed with Col-0, we detected the segregating T-DNA in the CLCf locus in approximately 33 % of individuals when *clcd clcf/+* was used as the female parent. However, when *clcd clcf/+* was used as pollen donor, the respective T-DNA was present in only 2 % of the offspring (Figure 3E). By introduction of the UBQ10:CLCf-mRFP construct we were able to rescue the *clcd clcf* double mutant. Nevertheless, plants were reduced in growth compared to wildtype indicating that either CLCf-mRFP is only partially functional or overexpression is detrimental (Supplementary Figure S2B).

**Figure 3:**
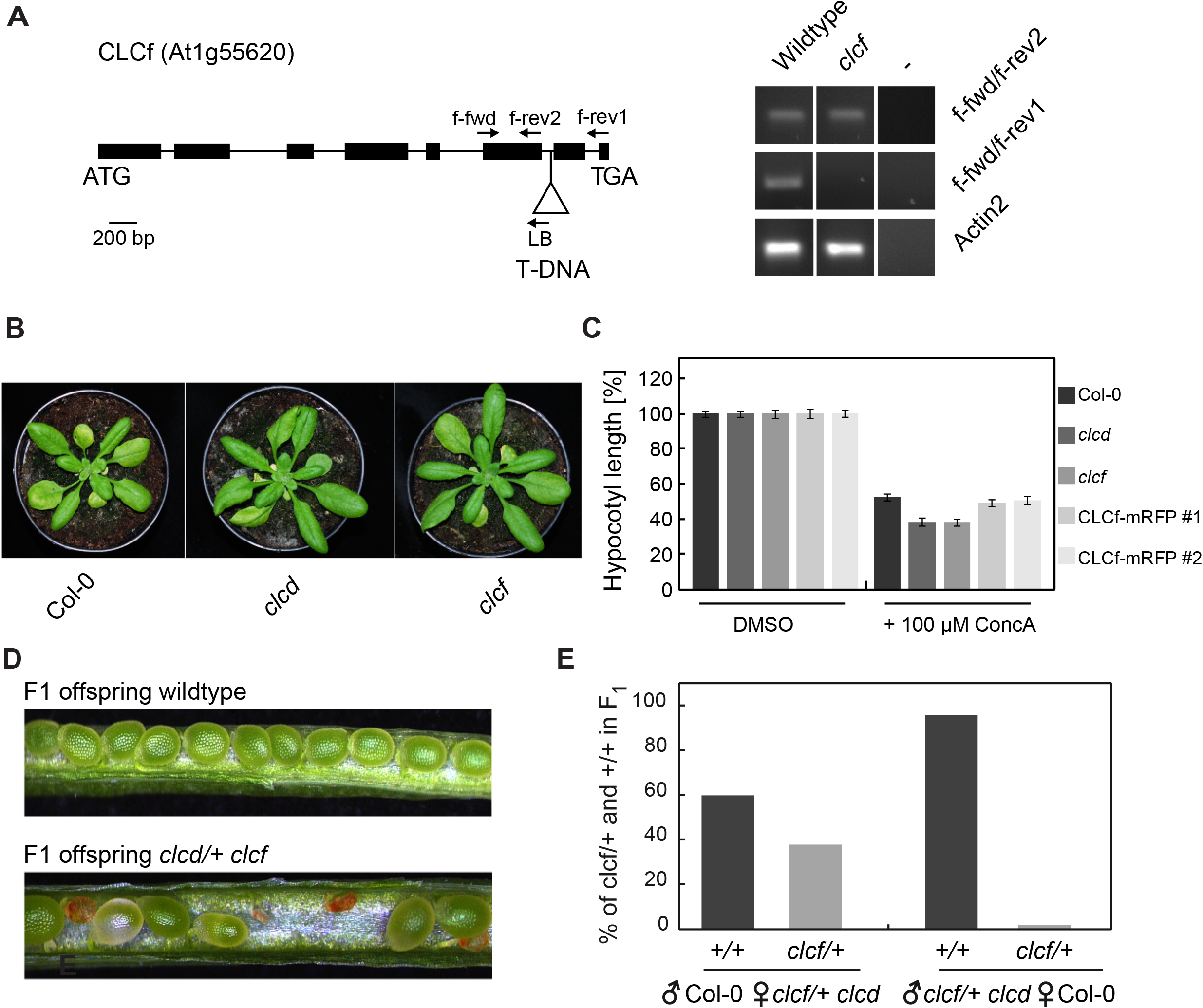
CLCd and CLC-f act redundantly. (**A**) Schematic overview of the CLCf gene. Exons are indicated as black boxes. Arrowheads indicate primers used for genotyping of the T-DNA insertion in CLCf (SALK_112962) using cDNA as a template. (**B**) Images of Col-0 wildtype, *clcd* and *clcf* 20 days after stratification. Plants have been cultivated under long day conditions (16h light/8h dark cycles). **(C)** Hypocotyl length measurements of 5-day-old etiolated seedlings grown on 1 % phytoagar, 5 mM MES-KOH (pH 5.8) supplied with 1 µM ConcA or DMSO. Representative data of one out of three independent experiments are shown. Error bars represent SE with n of 30. **(D)** Seed set of F1 offspring of wildtype Col-0 and *clcd/+ clcf*. Aborted or degenerated ovules indicate that the homozygous *clcd clcf* double mutant is lethal. **(E)** Presence of CLCf T-DNA-insertion or wildtype allele was analysed in the F1 generation of reciprocal crosses of *clcf clcd/+* and wildtype Col-0. For each cross n of 50 plants were analysed.

### CLC activity at the TGN/EE is required for cell expansion

To study the function of CLCd and CLCf during vegetative growth we employed inducible expression of an artificial microRNA (amiR) to knock-down *CLCf* in the *clcd* mutant background. An amiR targeting CLCf was designed and cloned under control of the 6xOP dexamethasone (Dex)-inducible promoter/LhGR system (Craft et al., 2005). The amiR construct (Figure 4A) was introduced into the *clcd* background expressing CLCf-mRFP and two independent homozygous lines were established. Induction of the amiR by 72 h treatment with 60 µM Dex reduced CLCf-mRFP protein levels to approximately 50 % *in clcd amiR-clcf #1* (Figure 4B) and less than 5 % *in clcd amiR-clcf #2* (Figure 4C). When an additional mCLCf-GFP construct in which the amiR recognition site was mutated (*mCLCf*) was introduced, no reduction in mCLCf-GFP levels was observed after DEX-induction indicating that amiR-expression is causative for the observed reductions in CLCf protein levels (Figure 4A, 4D). Upon Dex induction, root length was reduced to approximately 85 % in *clcd amiR-clcf #1* and to around 50 % in *clcd amiR-clcf #2* (Figure 4E, 4F). In line with these results, we observed stunted cells in the elongation- and differentiation zone as well as arrested root hair bulges in both induced *clcd amiR-clcf* lines (Figure 4E). Taken together, the function of TGN/EE-localised CLCs, like the TGN/EE-localised V-ATPase, is required for cell expansion.

**Figure 4:**
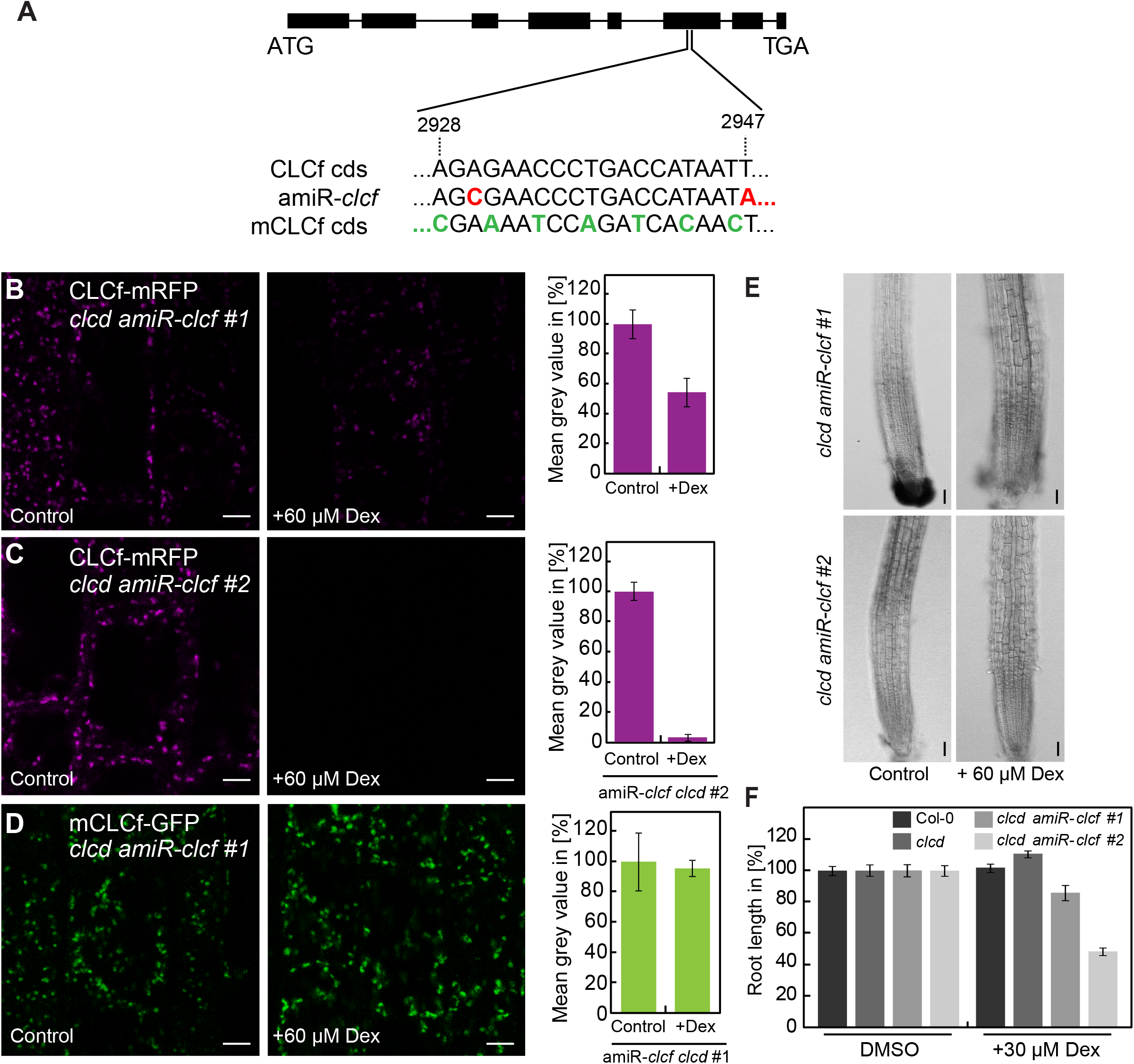
CLC-activity at the TGN/EE is required for cell expansion. **(A)** Schematic overview of the *CLCf* gene and the binding site for the artificial micro RNA targeting *CLCf*. Bases in red are altered in the amiR sequence compared to *CLCf*. Bases in green represent silent mutations introduced into the amiR binding site rendering *mCLCf* resistant.**(B-D)** Quantification of fluorescence was performed in cells of the root elongation zone of 7-day-old seedlings (**B)** mRFP fluorescence of UBQ10:CLCf-mRFP in *clcd amiR-clcf #1* after 72 h control or 60 µM Dex treatment. **(C)** mRFP fluorescence of UBQ10:CLCf-mRFP in *clcd amiR-clcf #2* after 72 h control or 60 µM Dex treatment. **(D)** GFP fluorescence of mCLCf-GFP in *clcd amiR-clcf #1* after 72 h control or 60 µM Dex treatment. **(B-D)** Scale bars represent 5 µm. Error bars represent SD of 3 independent experiments. For each experiment 15 seedlings were measured. **(E)** Phenotype of root tips of *clcd amiR-clcf #1* and *clcd amiR-clcf #2*, 72 h after transfer to 0.5 x MS medium, pH 5.8, 1 % phytoagar +/-60 µM Dex. Scale bar represent 50 µm. **(F)** Root length measurements of 7-day-old Arabidopsis seedlings grown on 0.5 x MS, pH 5.8, 1 % phytoagar. Plates were supplied with 30 µM Dex or equivalent amounts of DMSO. Error bars represent SE with n of 30 roots.

### Reduced CLC activity causes hyper-acidification of the Golgi stack

To measure pH in the TGN/EE of *clcd amiR-clcf #*2 we used an improved version of SYP61-pHusion in which we replaced mRFP against mCherry as it combines improved photostability and brightness with identical pH stable emission properties (Shaner et al., 2005). In vivo calibration of Arabidopsis lines expressing SYP61-pHusion2 showed that dynamic range and emission properties are comparable to the original version (Supplementary Figure S3). However, signals were less prone to photobleaching and less excitation energy was required. As expected, TGN/EE pH of *clcd* and *clcf* single mutants was found to be similar to wildtype (Figure 5B). To address if CLC-knock-down affects pH in the TGN/EE, we introduced UBQ10:SYP61-pHusion2 into the strong *clcd amiR-clcf* #2 knock-down line. In vivo pH measurements and calibrations were performed in all genetic backgrounds to exclude possible quenching effects of Cl^-^ on pHusion2 (Supplementary Figure 3A). Neither average pH (Figure 5C) nor the distribution of values for individual TGN/EEs (Figure 5D) were found to be affected in *clcd amiR-clcf* plants. However, we noted that the SYP61-pHusion2 signal in *clcd amiR-clcf* #2 appeared more diffuse after induction indicating that the sensor localization might be affected (Figure 5E). We thus next used ST-pHusion to measure pH in the neighboring trans-Golgi cisternae (Figure 5A) and found that it was significantly lower in *clcd amiR-clcf* #*2* after Dex-induction (Figure 5F, Supplementary Figure 3B).

**Figure 5:**
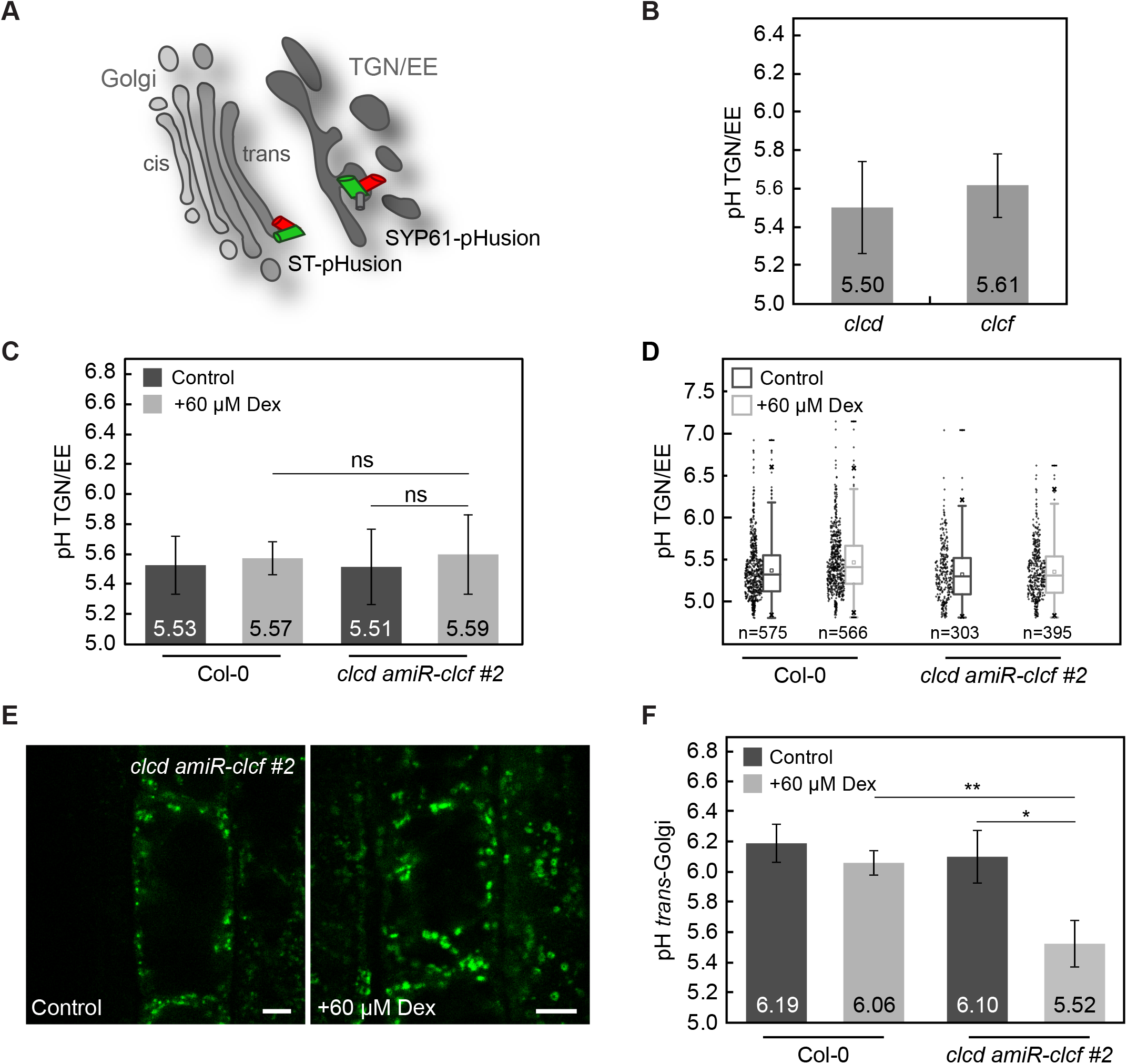
TGN/EE and *trans*-Golgi pH measurements. **(A)** Cartoon depicting the localization of the genetically encoded sensors ST-pHusion and SYP61-pHusion. **(B-F)** In vivo pH measurements have been performed cells of the root elongation zone of 7-day-old Arabidopsis seedlings stably expressing the respective pH sensors. **(B)** TGN/EE pH measurements in *clcd* and *clcf* expressing P16:SYP61-pHusion. **(C)** TGN/EE pH measurements in wildtype Col-0 and *clcd amiR-clcf #2* expressing UB10:SYP61-pHusion2 after 72 h control or DEX treatment. **(D)** Measurement of TGN/EE pH in individual particles positive for SYP61-pHusion2. Data were retrieved from 5 individual cells. **(E)** Signal distribution of UBQ10:SYP61-pHusion2 in cells of the root elongation zone in *clcd amiR-clcf #2* after 72 h control or Dex treatment. Scale bars represent 5 µm. **(F)** *Trans*-Golgi pH measurements in wildtype Col-0 and *clcd amiR-clcf #2* expressing UBQ10:ST-pHusion after 72 h control or Dex treatment. **(B-D, F)** Error bars represent SD of 3 independent experiments. For each experiment 15 seedlings were analysed. Significant differences are indicated by asterisks (ns indicates not significant, * indicates *p*-value < 0.05, ** indicates *p*-value < 0.01, Student’s t-test).

### Reduced CLC activity causes VHA-a1 to spread into the trans-Golgi

Hyper-acidification of the *trans*-Golgi cisternae could be caused by the accumulation of the V-ATPase as it transits on its way to the TGN/EE. Hence, we introduced VHA-a1-GFP (Dettmer et al., 2006) into *clcd amiR-clcf* and quantified colocalization with the endocytic tracer FM 4-64. Whereas colocalization of VHA-a1-GFP with FM4-64 did not change after 3 days Dex treatment in wildtype Col-0 (Figure 6A, 6B) TGN/EEs clustered and inflated in Dex-induced *clcd amiR-clcf* seedlings (Figure 6C-6E). Part of the VHA-a1-GFP signal was present in ring-like structures reminiscent of *trans*-Golgi signals (Figure 6D). Quantification based on Manders’ correlation coefficients confirmed a decreased signal overlap between VHA-a1-GFP and FM 4-64 (M1) in Dex-induced but also in uninduced *clcd amiR-clcf* seedlings. In line, Manders’ 2 (M2) correlation coefficients, which represent the fraction of FM 4-64 signal overlapping with VHA-a1-GFP, were already strongly increased in *clcd* and were even further elevated in Dex-induced *clcd amiR-clcf* seedlings (Figure 6F). In summary, CLC function is required to maintain the functional boundary between trans-Golgi and TGN/EE and could therefore affect both secretory and endocytic trafficking.

**Figure 6:**
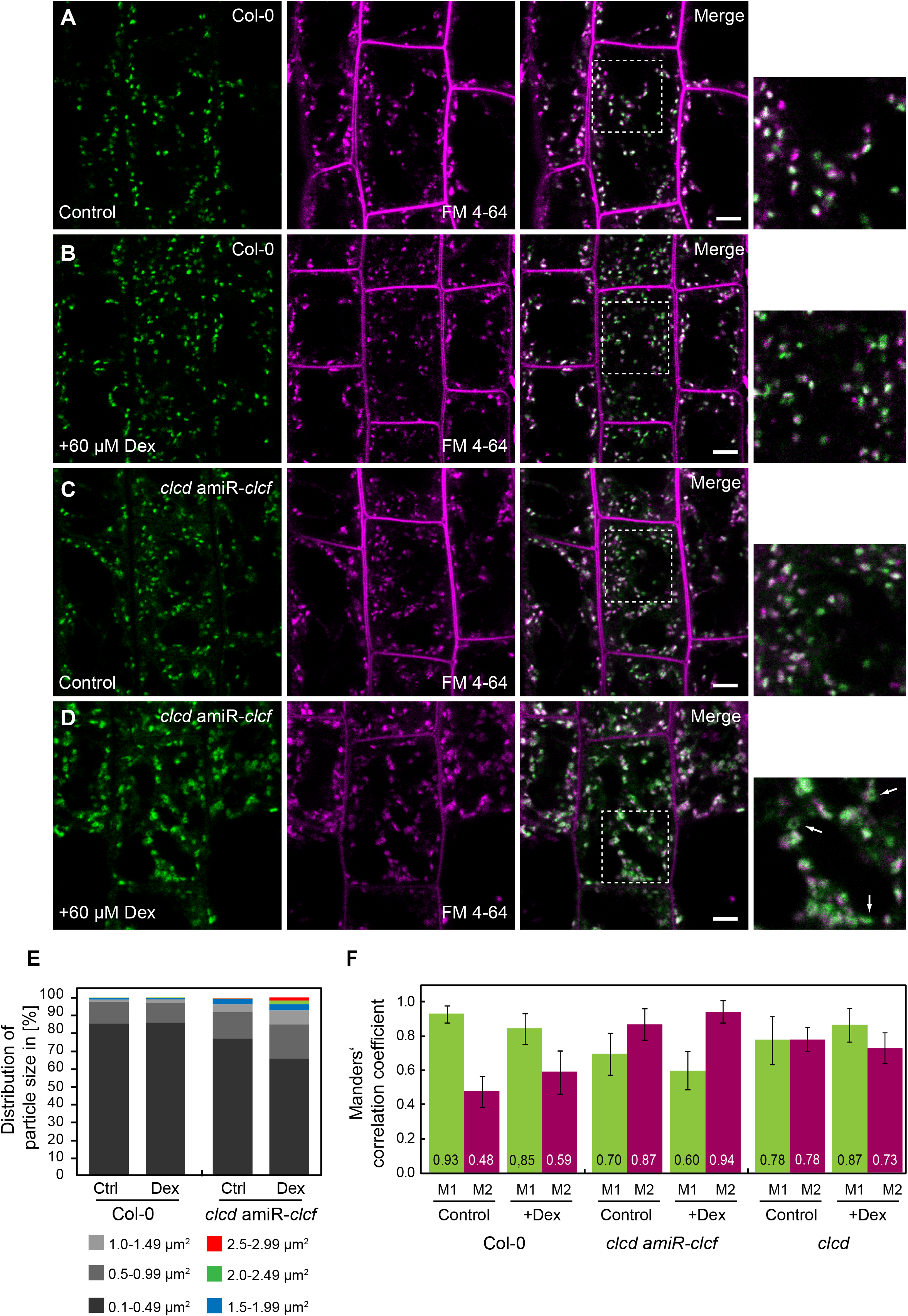
VHA-a1 is partially mislocalized to the Golgi stack in *clcd amiR-clcf*. **(A-D)** Localisation analyses was performed in cells of the root elongation zone of 7-day-old Arabidopsis seedlings stably expressing VHA-a1-GFP (green). Seedlings were stained with 1 µM FM 4-64 for 15 min (magenta). Seedlings were imaged 72 h after transfer to 0.5 x MS medium, pH 5.8, 1 % phytoagar +/-60 µM Dex. Dashed line marks magnified area. Scale bars represent 5 µm. **(A, B)** Wildtype Col-0. **(C, D)** *clcd amiR-clcf*. **(D)** Arrowheads indicate ring-like structures resembling signals for the *trans*-Golgi stack. **(E)** Particle size analyses was performed in cells of the root elongation zone of 7-day-old Arabidopsis seedlings stably expressing SYP61-pHusion2. Seedlings were analysed 72 h after transfer to 0.5 x MS medium, pH 5.8, 1 % phytoagar +/-60 µM Dex. 15 seedlings for each genotype and treatment were analysed. **(F)** Colocalization analyses has been performed on samples depicted in **(A-D)**. Manders’ correlation coefficients M1 and M2 have been calculated, where M1 represents the fraction of VHA-a1-GFP overlapping with FM4-64 and M2 represents the fraction of FM4-64 overlapping with VHA-a1-GFP. Data were acquired from 10 cells of 10 individual roots. Error bars represent SD.

### Reduced CLC activity has no major effect on post-Golgi trafficking

To assess if reduced CLC activity affects post-Golgi trafficking, we crossed plants expressing a secreted version of mRFP (secRFP; Batoko et al., 2000; Zheng et al., 2005) or the plasma membrane-localized auxin efflux carrier PIN1 (PIN-FORMED1)-GFP (Benková et al., 2003) with both inducible *clcd amiR-clcf* lines. Both markers were imaged 72 h after Dex-induced amiR expression in homozygous F2 lines. However, neither intracellular accumulation of secRFP (Figure 7A, 7C, 7E) or mislocalization of PIN1-GFP (Figure 7B, 7D, 7F) were observed in *clcd amiR-clcf* after induction. Furthermore, we assessed if post-Golgi trafficking to the vacuole is affected by a FM 4-64 chase experiment. We detected tonoplast labelling in wildtype Col-0 and in *clcd amiR-clcf* lines after 100 min FM 4-64 uptake (Figure 7G-J). In summary, we could not detect major effects on secretory and vacuolar trafficking in *clcd amiR-clcf* and it thus remains to be determined how the knock-down of CLC function in the TGN/EE leads to the observed inhibition of cell expansion.

**Figure 7:**
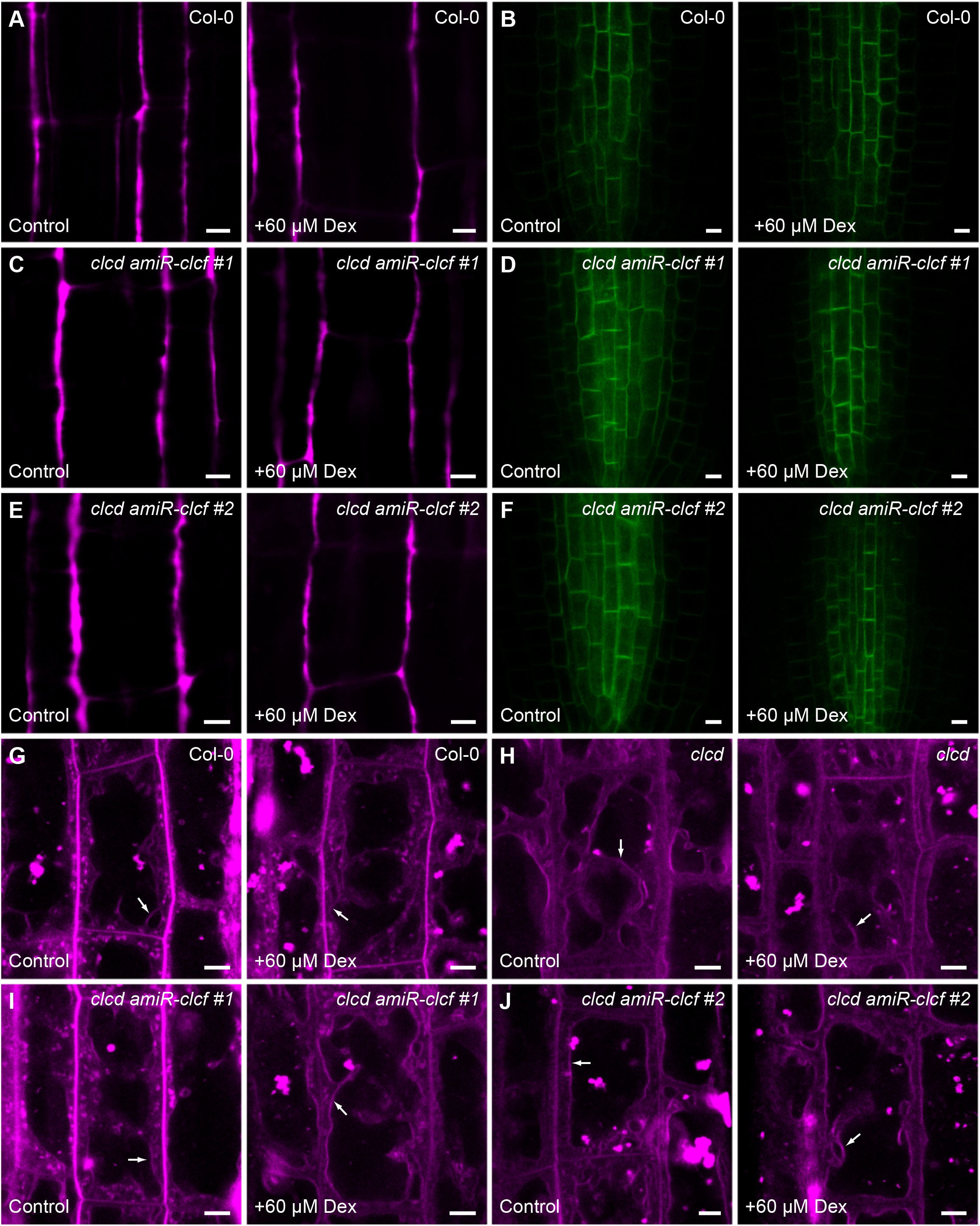
Reduced CLC activity has no major effect on post-Golgi trafficking. Imaging was performed in root cells of 7-day-old Arabidopsis seedlings 72 h after transfer to 0.5 x MS medium, pH 5.8, 1 % phytoagar +/-60 µM Dex. **(A, C, E)** secRFP signal distribution in wildtype Col-0 **(A)** in *clcd amiR-clcf #1* **(C)** and in *clcd amiR-clcf #2* **(E)** under control and Dex-induced conditions. **(B, D, F)** PIN1-GFP signal distribution in wildtype Col-0 **(B)** in *clcd amiR-clcf #1* **(D)** and in *clcd amiR-clcf #2* **(F)** under control and Dex-induced conditions. **(G-J)** Uptake and distribution of FM4-64 after 100 min of staining with 1 µM in wildtype Col-0 **(G)**, in *clcd* **(H)**, in *clcd amiR-clcf #1* **(I)** and in *clcd amiR-clcf #2* **(J)** under control and Dex-induced conditions. Arrowheads indicate tonoplast. **(A-J)** Scale bars represent 5 µm.

## Discussion

The TGN/EE is a highly dynamic compartment that many proteins pass en route to their destination. Among the residents that stay behind proteins that maintain pH- and ion homeostasis are dominant. As the primary proton-pump the V-ATPase plays a central role, however the presence of several other transport proteins indicates that their concerted action is required (Schumacher, 2014; Sze and Chanroj, 2018). NHX5 and NHX6, two K^+^/H^+^-antiporters reside in the TGN/EE and in MVBs, however it remains to be determined if their function as proton leaks to alkalise the endosomal lumen during maturation or their effect on luminal potassium concentration is responsible for the severe phenotype of *nhx5 nhx6* mutants (Bassil et al., 2011; Dragwidge et al., 2019). Similarly, CLCd is localised at the TGN/EE and could contribute to the shunt conductance required to balance the electrogenicity of the V-ATPase or maintain the concentration of chloride or other anions in the lumen of the TGN/EE (von der Fecht-Bartenbach et al., 2007). However, compared to mutants with reduced V-ATPase activity (Brüx et al., 2008; Schumacher et al., 1999) the *clcd* mutant has a rather mild phenotype and we thus reasoned that it might act redundantly with CLCf, the only member of the Arabidopsis CLC-family that remained largely uncharacterised. The evolutionary origin of all CLCs can be traced back to prokaryotes but CLCe and CLCf belong to a particular subgroup that is closely related to cyanobacterial CLCs indicating that they might be of plastidial rather than mitochondrial origin. Indeed, subgroup II is confined to the green lineage and CLCe is found in thylakoid membranes (Marmagne et al., 2007; Monachello et al., 2009). Members of the CLCf clade are characterised by the presence of an N-terminal extension of variable length and low sequence conservation that does not contain features predictive of chloroplast localization. Based on biochemical approaches CLCf has been reported to be localised in the chloroplast outer envelope (Teardo et al., 2005). However, based on transient expression of a GFP-fusion protein CLCf has been localised to Golgi membranes and the fact that CLCf rescues the *gef1* mutant that lacks the Golgi-localized sole yeast CLC independently confirms that CLCf is not a plastidial protein (Marmagne et al., 2007). We have shown here that CLCf colocalizes with CLCd and VHA-a1 in the TGN/EE and it will be of interest to determine if localisation of the two only distantly related CLCs is mediated by a similar mechanism. Not only is knowledge regarding TGN/EE-retention or retrieval of integral membrane proteins limiting, functional analysis of CLCd and CLCf will require their rerouting to membranes amenable to patch-clamp analysis such as the tonoplast. Our attempts at rerouting have so far not been successful and we thus have to rely on predictions based on homology modelling. Whereas CLCd belongs to the well characterised subgroup I and has all the hallmarks of a H^+^-coupled anion transporter with a strong preference for Cl^-^ over NO_3-_, CLCf has been suggested to have channel-like properties as the characteristic proton glutamate seemed to be missing (Zifarelli and Pusch, 2010). Based on homology modelling we have shown here that E300, a residue conserved in all subgroup II CLCs could occupy the same position as E312, the proton glutamate of chicken CLC-7 (Schrecker et al., 2020) indicating that CLCf and members of subgroup II in general are proton-coupled antiporters. Despite the absence of experimental evidence regarding selectivity, chloride seems to be the most likely substrate yet other anions can of course not be excluded. In any case, the results of our genetic analysis are in line with CLCd and CLCf having redundant functions as H^+^/Cl^-^-antiporters in the TGN/EE. Both single mutants can be distinguished from wildtype based on their increased sensitivity to the V-ATPase inhibitor ConcA, a phenotype that could be indicative of their role as shunt conductances that counter the charge imbalance created by H^+^-pumping (Sze and Chanroj, 2018). CLCd has been reported to negatively regulate pathogen-triggered immunity in response to flg22 (Guo et al., 2014) whereas reduced V-ATPase activity dampens the flg22 response (Keinath et al., 2010) indicating that the functional relation between CLCs and the V-ATPase might be more complex and a more detailed characterisation of the single mutants might be required to reveal non-redundant functions. However, the fact that the combined loss of CLCd and CLCf causes male gametophyte lethality, a phenotype also observed in V-ATPase null mutants (Dettmer et al., 2005; Lupanga et al., 2020) also supports a role for CLC-activity in supporting acidification of the TGN/EE. Further analysis is required to determine if the underlying pollen defects are indeed similar, yet developing pollen is not the most suitable experimental system to study CLC-function in the TGN/EE in more detail. To investigate the role of CLCs in TGN/EE-acidification, we established an inducible amiRNA-based knock-down system for *CLCf* in a *clcd* null background and used an improved version of theTGN-localised pH-sensor SYP61-pHusion to measure pH. To our surprise, reduced CLC activity did not affect pH in the TGN/EE but instead caused an increased acidification of *trans*-Golgi cisternae. It remains to be determined if impaired cell elongation is the consequence of improper TGN/EE anion homeostasis or if the altered *trans*-Golgi pH is responsible.

Although these results do not seem to support a role for CLCs as shunt conductances, it needs to be taken into account that the knock-down is likely incomplete so that residual CLC activity might still be sufficient to maintain V-ATPase activity in the TGN/EE. Hyper-acidification of the *trans*-Golgi could affect different aspects of cell wall biosynthesis. Cellulose synthase complexes (CSCs) are assembled in the Golgi stack (Zhang et al., 2016) and suboptimal pH could affect CSC assembly or structure of individual CesA subunits. The major pectin component hemigalacturonan (HG) is synthesized in the Golgi apparatus and methyl-esterification occurs in the *trans*-Golgi. The more acidic *trans*-Golgi pH could alter pectin methyltransferase activity leading to overall lower HG methyl-esterification and it been suggested that enzymes responsible for pectin stability and degradation are co-processed with its substrate in the Golgi stack (Anderson, 2016). However, less obvious connections also need to be taken into account. In yeast mutants lacking Gef1, the only CLC protein, the multicopper oxidase Fet3 fails to acquire copper ion cofactors and by the introduction of an uncoupling mutation that allows acidification but abolishes antiport activity it was shown that Gef1 has marked effects on cellular glutathione homeostasis (Braun et al., 2010). Also in mammalian cells, uncoupling mutations that convert CLCs from antiporters to channels have revealed that the capacity for luminal chloride accumulation is a critical function of CLC-proteins (Novarino et al., 2010; Weinert et al., 2020) and it has been suggested that endomembrane compartments exploit Cl^-^ for volume regulation (Stauber and Jentsch, 2013). We still know very little about the spatio-temporal aspects of TGN/EE-maturation and its functional separation from the Golgi stack. However, the effects of the monovalent cation ionophore monensin that induces rapid osmotic swelling of the TGN/EE followed by trans-Golgi cisternae (Dragwidge et al., 2019; Zhang et al., 1993) underline that volume regulation is a critical aspect. Recently, the Cation Chloride Cotransporter 1 (CCC1) of Arabidopsis has been shown to be localized at the TGN/EE and to affect pH and ion homeostasis. By mediating both cation and anion efflux CCC1 complements the ion transport circuit of the TGN/EE (McKay et al., 2020). However, the circuit might not yet be complete. In mammalian cells, the proton-activated Cl^−^ (PAC) channel releases Cl^-^ from the endosomal lumen but in contrast to CCC1 deletion of PAC leads to hyper-acidification of endosomes (Osei-Owusu et al., 2021; Ullrich et al., 2019). It remains to be determined if a PAC1-like protein is present in plants but in any case determining the functional dependencies between the different players is likely to provide further insight into the physiology of the TGN/EE.

## Acknowledgements

We would like to thank Fabian Fink for technical assistance and Beate Schöfer for plant care. This work was supported by the Deutsche Forschungsgemeinschaft within SFB1101 (A02) to KS and KR4675/2-1 to MK.

## Author contributions

Stefan Scholl, Conceptualization, Investigation, Methodology, Writing - original draft; Stefan Hillmer, investigation Melanie Krebs, Writing – review and editing, Funding acquisition; Karin Schumacher, Conceptualization, Resources, Supervision, Funding acquisition, Investigation, Methodology, Writing - original draft, Project administration, Writing -original draft, Writing - review and editing

## Materials and methods

### Phylogenetic analysis and homology modelling

#### Phylogenetic analyses

CLC protein sequences (Supplementary Table S1) were pairwise aligned (Global alignment with free end gaps with gap open penalty and gap extension penalty of 3) using the Blosum 45 cost matrix and a neighbor-joining tree was built using the Jukes-Cantor genetic distance model in Geneious 10.1.3 (Biomatters Ltd., Auckland, New Zealand).

#### Homology modeling

The annotated, full length amino acid sequence for CLC was submitted as a query without reference structure to the online protein structure prediction tool I-Tasser (Yang and Zhang, 2015). The top ranking model among the structural analogs in PDB was identified to be 7JM6A (CLC-7) with a TM-score of 0.772. Images of the resulting structure were generated using ChimeraX-1.1.1 (Pettersen et al., 2021).

### Cloning procedures

To clone *UBQ10:CLCf-mRFP*, the full length coding sequence (CDS) of *CLCf (At1g55620)* was amplified without stop codon from *Arabidopsis thaliana* Col-0 cDNA using primers CLCf-*Aat*II-Fw and CLCf-noSTOP-*Pvu*I-Rv. The CDS of mRFP was amplified from SYP61-pHusion (Luo et al. 2015) using mRFP-*Pvu*I-Fw and mRFP-*SalI*-Rv. Both fragments were subcloned into pJET1.2blunt (ThermoFisher Scientific, MA, USA) and sequenced. The respective fragments were released from pJET1.2blunt using restriction enzymes *Aat*II, *Pvu*I and *Sal*I, respectively and ligated with *pUTKan* (Krebs et al., 2012) that has been digested with *Aat*II and *Sal*I to finally receive *UBQ10:CLCf-mRFP*.

*UBQ10:ST-GFP* construct was generated and assembled using the GreenGate (GG) cloning system (Lampropoulos et al., 2013). Modules used to generate *UBQ10:ST-GFP* via GG reaction are listed in Supplementary Table S2.

*UBQ10:CLCd-GFP* was generated using GG cloning. To generate the entry module *pGGC-CLCd*, two internal *Eco31*I F sites were removed from the wildtype *CLCd* sequence by introduction of silent mutations. Furthermore, the entire sequence of *CLCd* intron 1 was added due to toxic effects of the gene product for bacteria. For that purpose, four separate PCR reactions were performed. First, a 86-bp fragment of the 5’
sregion of *CLCd* was amplified from cDNA with primers GG-CLCd-C-fwd and GG-CLCd-Int1-rev thereby mutating nucleotide at position 54 from T to A, followed by a second PCR, in which a 518-bp fragment was amplified from Arabidopsis genomic DNA with primers GG-CLCd-Int2-fwd and GG-CLCd-Intron1-rev. In a third PCR reaction, using primers GG-CLCd-Intron1-fwd and GG-CLCd-Int2-rev, a 881-bp fragment was amplified, thereby mutating A to G at position 1731. The last PCR reaction using primers CLCd-Int3-fwd and CLCd-noSTOP-rev amplified a 677-bp product of the *CLCd* CDS. All four PCR products were digested with *Eco31*I and ligated into *Eco31*I digested *pGGC000 (Lampropoulos et al*., *2013)* to receive *pGGC-CLCd*. A correct clone was verified by sequencing. *pGGC-CLCd* was then used in a GG reaction with modules listed in Supplementary Table S2 to generate *UBQ10:CLCd-GFP*.

The *Dex:amiR-clcf* construct was generated using GG cloning. An artificial microRNA (amiR) against *CLCf* (5’-AGCGAACCCTGACCATAATA-3’) was designed using the WMD3 microRNA design tool (Schwab et al., 2006). The amiR targets the *CLCf* CDS between position 2928 and 2947 and was introduced into *pRS300* (kindly provided by Detlef Weigel) using oligonucleotides amiR-CLCf-I, amiR-CLCf-II, amiR-CLCf-III and amiR-CLCf-IV according to protocols provided on www.weigelworld.org. The 439-bp amiR cassette was amplified using primers GG-amiR-FW-B-overhang-*EcoR*I and GG-amiR-RV-C-overhang-*BamH*I digested with *Eco31*I, and integrated into *pGGI000 (Lampropoulos et al*., *2013)* to receive p*GGI-amiR-clcf*. The correct *amiR-clcf* cassette was verified by sequencing and subsequently used to generate the intermediate vector *pGGN-6xOP:amiR-clcf* via GG reaction with modules listed in Supplementary Table S2. Afterwards a final GG reaction was performed with *pGGN-6xOP:amiR-clcf, pGGM-UBQ10:Lh4GR* (Lupanga et al., 2020) and the destination vector *pGGZ003* (Lampropoulos et al., 2013) to receive *Dex:amiR-clcf*.

*VHP1:mCLCf-GFP* was created using GG cloning. To generate an amiR-resistant version of *CLCf* (*mCLCf*) several bases within the 20 bp amiR recognition site were mutated without changing the amino acid sequence (5’-CGAAAATCCAGATCACAACT-3’). For this, the full length *CLCf* CDS without stop codon was amplified from *UBQ10:CLCf-mRFP* in two separate PCR reactions using primers GG-CLCf-C-fwd and GG-mCLCf-rev and mCLCf-fwd and GG-CLCf-C-rev which resulted in 1814-bp and 572-bp fragments, respectively. Both fragments were *Eco31*I digested and inserted into *pGGC000* to receive *pGGC-mCLCf*. A correct clone was confirmed by sequencing. *pGGC-mCLCf* was then used in a GG with modules listed in Supplementary Table S2 to receive *VHP1:mCLCf-GFP*.

For *UBQ10:SYP61-pHusion2*, the bleaching-sensitive mRFP moiety of pHusion has been exchanged by the more bleaching-resistant mCherry. For this, the CDS of mCherry was amplified from GCaMP6f-mCherry (Waadt et al., 2017)with primers mCherry-*Hind*III-fwd and mCherry-noSTOP-*Eco31*I-rev introducing *Hind*III sites at the 5’ and a unique 4 bp overhang (TAAC) at the 3’end. EGFP was amplified from *P16:SYP61-pHusion* (Luo et al. 2015) using primers EGFP-*Eco31*I-fwd and pHusion-*Sal*I-rv which span the full length CDS of EGFP plus a 5 aa linker (AVNAS). A fragment encoding *SYP61* was released with *BamH*I and *Hind*III from *P16:SYP61-pHusion* (Luo et al. 2015). The three fragments obtained, *SYP61* (*BamH*I/*Hind*II), *mCherry* (*Hind*III/TAAC) and *EGFP* (TAAC/*Sal*I) were ligated into *BamH*I/*Sal*I-digested *pUTkan* ((Krebs et al., 2012)) to receive *UBQ10:SYP61-pHusion2*.

For *UBQ10:ST-pHusion* the CDS of rat sialyltransferase (ST) was amplified from *35S:ST-pHluorin* (Martinière et al., 2013) using primers ST-*Aat*II-fwd and ST-noSTOP-*Pvu*I-rev which removed the stop codon. The CDS of pHusion was amplified from *P16:SYP61-pHusion* with primers pHusion-*Pvu*I-fwd and pHusion-*Sal*I-rev. The ST and pHusion PCR fragments were subcloned into pJETblunt1.2. Correct clones were verified by sequencing. *ST* and *pHusion* fragments were released from their respective pJET backbones using *Aat*II/*Pvu*I and *Pvu*I/*Sal*I, respectively and finally ligated into the *Aat*II/*Sal*I digested backbone of the binary vector *pUTBar. pUTBar* contains a multiple cloning site that is flanked by the *UBQ10* promoter from Arabidopsis and the *RBCS* terminator from pea and it confers Basta resistance to plants. *pUTBar* is a descendant of *pTBar* which is a derivative of *pPZP312* that originated from the *pPZP* vector series (Hajdukiewicz et al., 1994).

All primers used for cloning of the constructs are listed in Supplementary Table S3.

### Plant materials and growth conditions

If not stated otherwise, *Arabidopsis thaliana* ecotype Col-0 seeds were grown on media containing 0.5 x MS, 0.5 % sucrose with pH adjusted to 5.8 with KOH. Plates were solidified using 0.6 % phytoagar. Agar and MS basal salt mixture were purchased from Duchefa (Duchefa Biochemie, Haarlem, Netherlands). Seeds were surface-sterilized with EtOH followed by stratification for 48 h at 4 °C. Seedlings were grown at 22 °C with cycles of 16 h light and 8 h darkness.

### Hypocotyl and root length measurements

For hypocotyl measurements, seeds were surface-sterilized and plated on medium consisting of 1% phytoagar; 5 mM MES-KOH, pH 5.8. After stratification at 4 °C for 48 h seeds were exposed to 5 h of light (120 μmol m^-2^ s^-1^) before being wrapped in a double layer of aluminium foil. Seedlings were then grown in the dark for 4 days at 22 °C. To measure hypocotyl length, seedlings were sandwiched between two layers of acetate and scanned. The digitized images were analyzed using the segmented line tool in Fiji (Schindelin et al. 2012). For the inhibitor treatment, 100 nM ConcanamycinA (Santa Cruz Biotechnology, TX, USA) or the equivalent volume of DMSO was added to the plant growth medium. For each condition, hypocotyls from 30 seedlings were measured.

For root length measurements, seedlings were grown for 7 days on vertical plates containing 0.5 x MS, pH 5.8, 1 % phytoagar at long day conditions. Plates were supplied with 30 μM Dexamethasone (Dex; D4902, Sigma-Aldrich, MO, USA) or equivalent amounts of DMSO.To measure root length, seedlings were sandwiched between two layers of acetate and scanned. The digitized images were analyzed using the segmented line tool in Fiji (Schindelin et al. 2012).

### Seed set analyses

Mature siliques of *Arabidopsis thaliana* ecotype Col-0 wildtype and the *clcd/+ clcf* mutant, harbouring one copy of the T-DNA insertion for CLCd (*clcd-1*) and two copies of the T-DNA insertion for CLCf (*clcf-1*), grown under long day conditions, were opened along the seed valve from base to tip. Phenotype of the seed set was documented using a Zeiss SteREO Discovery V20 (Carl Zeiss Microscopy GmbH, Oberkochen, Germany).

### Reciprocal crosses

To analyse whether transmission through either male or female gametes is dependent on *CLCf*, reciprocal crosses between wildtype Col-0 and *clcd/+ clcf* were performed. The resulting F1 offspring was analysed for presence of CLCf wildtype and *clcf-1* T-DNA insertion allele via genotyping as stated above. 50 plants were analysed for each reciprocal cross.

### Stable transformation of Arabidopsis plants

*UBQ10:CLCf-mRFP* and *UBQ10:SYP61-pHusion2* were introduced into *Agrobacterium tumefaciens* strain GV3101:pMP90. *UBQ10:ST-GFP, UBQ10:CLCd-GFP and VHP1:mCLCf-GFP* were introduced into *Agrobacterium tumefaciens* strain ASE containing the *pSOUP* helper plasmid. *Arabidopsis thalian*a ecotype Col-0 wild-type plants were used for transformation via floral dip using standard procedures (Hellens et al., 2000). Transgenic lines were isolated on plates containing 0.5 x MS, 0.5 % sucrose, pH 5.8 (KOH) and 0.55 % phytoagar supplemented either with 50 µg/ml Kanamycin, 10 mg/ml BASTA, 11.25 µg/ml Sulfadiazine or 25 µg/ml HygromycinB.

### Isolation of a clcf T-DNA insertion allele

To study the function of CLCf, a homozygous T-DNA insertion allele (*clcf-1*) from the SALK collection (SALK_112962) was characterised. The position of the T-DNA insertion in intron 6 was verified by PCR using primers listed in Supplementary Table S3 followed by confirmation of the insertion site by sequencing. Gene-specific primers CLCf-fwd and CLCf-rev2 were used to detect exon 6 which is situated before the T-DNA insertion. A gene-specific primer pair (CLCf-fwd, CLCf-rev1) spanning the T-DNA insertion and a gene-/T-DNA-specific primer pair (CLCf-fwd, TL1-LB) was used to detect the T-DNA insertion in the *CLCf* locus. PCR amplification of an *ACTIN2* fragment with gene-specific primers was used as a control.

### Transgenic lines used in this study

Stable transgenic Arabidopsis lines expressing VHA-a1-GFP (Dettmer et al. 2006), VHA-a1-mRFP (von der Fecht-Bartenbach et al., 2007) P16:SYP61-pHusion (Luo et al. 2015), secRFP (Zheng et al., 2005) and PIN1-GFP (Benková et al., 2003) as well as the T-DNA insertion allele for *CLCd* (*clcd-1*; von der Fecht-Bartenbach et al. 2007) have been described previously. Transgenic plants expressing a combination of fluorescent protein-tagged proteins were generated by cross-pollination or transformation.

### Confocal laser scanning microscopy and image processing

Confocal laser scanning microscopy (CLSM) of Arabidopsis roots was performed with a Leica SP5II system equipped with an inverted DMI6000 microscope stand (Leica Microsystems GmbH, Wetzlar, Germany). Images were recorded using a HCX PL APO lambda blue 63.0×1.20 WATER UV or a HCX PL APO CS 20.0×0,70 IMM UV objective (Leica Microsystems GmbH, Wetzlar, Germany). GFP was excited with 488 nm, mRFP and FM4-64 were excited with 561 nm. Fluorescence was detected between 500-545 for GFP, between 620-670 nm for mRFP and between 670-750 nm for FM4-64 using hybrid detectors (HyD; Leica Microsystems GmbH, Wetzlar, Germany). Images for non-quantitative purposes were adjusted in brightness and contrast using Fiji (Schindelin et al., 2012).

### Quantification of average fluorescence intensities

Average fluorescence intensities were determined in roots of 7-day-old Arabidopsis seedlings expressing UBQ10:CLCf-mRFP or VHP1:mCLCf-GFP in the *clcd amiR-clcf* background 72 h after *amiR-clcf* induction with 60 μM Dex. Plants were pre-grown for four days on 0.5 x MS; pH 5.8; 0.55 % phytoagar and afterwards transferred to identical control medium or medium supplied with 60 μM Dex. Plants were further grown for 3 days at identical light conditions before imaging was performed. Imaging of mRFP and GFP was performed as described above. Average fluorescence intensity values were quantified with Fiji using a self-written macro (Appendix 1 S4). Per genotype and condition 15 images were taken. The experiment was repeated three times.

### Colocalization analyses

For colocalization analysis images were acquired using the HCX PL APO lambda blue 63.0 x 1.20 water immersion objective.The imaging parameters were as follows: 1024 x 1024 pixel resolution with a zoom of 2.5 or 512 x 512 pixel resolution with a zoom of 5.3 was chosen to adjust the voxel size to 96 nm. All microscope settings remained identical for each sample. All images were processed with Fiji (Schindelin et al., 2012) using a Gaussian Blur Filter with a sigma radius of 1. Pearsons’ and Spearmans’ correlation coefficients as well as scatter plots were calculated using the *PSC Colocalization* plugin (French et al., 2008) with a threshold level of 10. Manders’ overlap coefficient between VHA-a1-GFP and FM 4-64 positive endosomes was determined in 7-day-old Arabidopsis seedlings stably expressing VHA-a1-GFP in wildtype Col-0 or in a*miR-clcf clcd*. Prior to induction seedlings were grown for 4 days on 0.5 x MS; pH 5.8; 0.55 % phytoagar followed by transfer to identical control medium or medium supplied with 60 μM Dex. Seedlings were imaged 72 h after Dex induction and after 15 min of staining with 1 µM FM4-64 (T13320, ThermoFisher Scientific, MA, USA) in liquid medium (0.5 x MS medium, pH 5.8 with KOH).

To determine Manders’ overlap coefficient, images of 10 cells from 8-10 different seedlings within the root elongation zone were acquired using the HCX PL APO lambda blue 63.0 x 1.20 water immersion objective.The imaging parameters were as follows: 512 x 512 pixel resolution with a zoom of 5.3 was chosen to adjust the voxel size to 96 nm. All microscope settings remained identical for each sample. All images were processed with Fiji (Schindelin et al., 2012) using a Gaussian Blur Filter with a sigma radius of 1. Regions of interests were drawn manually to exclude the plasma membrane signal of FM4-64. M1 and M2 Manders’ overlap coefficients were calculated using the JACoP plugin in Fiji (Bolte and Cordelières, 2006) with a threshold of 40 for both channels.

### Particle size measurement of TGN/EEs

Surface area measurements of TGN/EEs and TGN/EE clusters were performed in root epidermal cells of 7-day-old Arabidopsis seedlings stably expressing UBQ10:SYP61-pHusion2 in wildtype Col-0 and in the *clcd amiR-clcf* background. Plants were pre-grown for 4 days and then transferred to identical control medium or medium supplemented with 60 μM Dex. Plants were grown for 3 days further on the respective medium prior to imaging. Images were acquired using the HCX PL APO lambda blue 63.0 x 1.20 water immersion objective at 512 x 512 pixel resolution with a pinhole size of 0.75 AU. The fluorescence signals of the GFP channel were used for size calculation in Fiji (Schindelin et al., 2012). Average background values were determined and subtracted from each image. Afterwards the Gaussian Blur Filter with a sigma radius of 1 was applied and images were thresholded (30/255). The particle size of single or clusters of TGN/EEs was determined using the Particle analyzer with particle sizes ranging from 0.1-3.0 μm^2^ and with a circularity of 0.05-1.00. Values were grouped according to area.

### pH-measurements in the TGN/EE and in the trans-Golgi

pH-measurements in the TGN/EE using P16:SYP61-pHusion, UBQ10:SYP61-pHusion2 and UBQ10:ST-pHusion were performed as described previously (Luo et al., 2015).

### High-Pressure Freezing, Freeze Substitution, and Electronmicroscopy

Entire root tips of 6-day-old Arabidopsis seedlings stably expressing *UBQ10:CLCf-mRFP* were harvested and processed as previously described (Scheuring et al., 2011). Freeze substitution was performed in a Leica EM AFS2 freeze substitution unit (Leica Microsystems GmbH, Wetzlar, Germany) in dry acetone supplemented with 0.3 % uranyl acetate as previously described (Hillmer et al., 2012). Ultrathin sections of 80 – 90 nm were obtained using a Leica Ultracut S (Leica Microsystems GmbH, Wetzlar, Germany) microtome with a diamond knife (Diatome Ltd, Nidau, Switzerland). Grid-mounted sections were incubated with an α-DsRed antibody in 1:200 dilution followed by incubation with a goat anti-rabbit (1:50) antibody linked to 10 nm colloidal gold particles. Immunolabeled sections were observed with a JEOL JEM-1400 (JEOL, Tokio, Japan) electron microscope operating at 80 kV. Micrographs were taken with a TVIPS FastScan F214 digital camera (TVIPS GmbH, Gauting, Germany). Brightness and contrast were adjusted using Fiji (Schindelin et al., 2012).

---

**Supplementary Figure S1:**
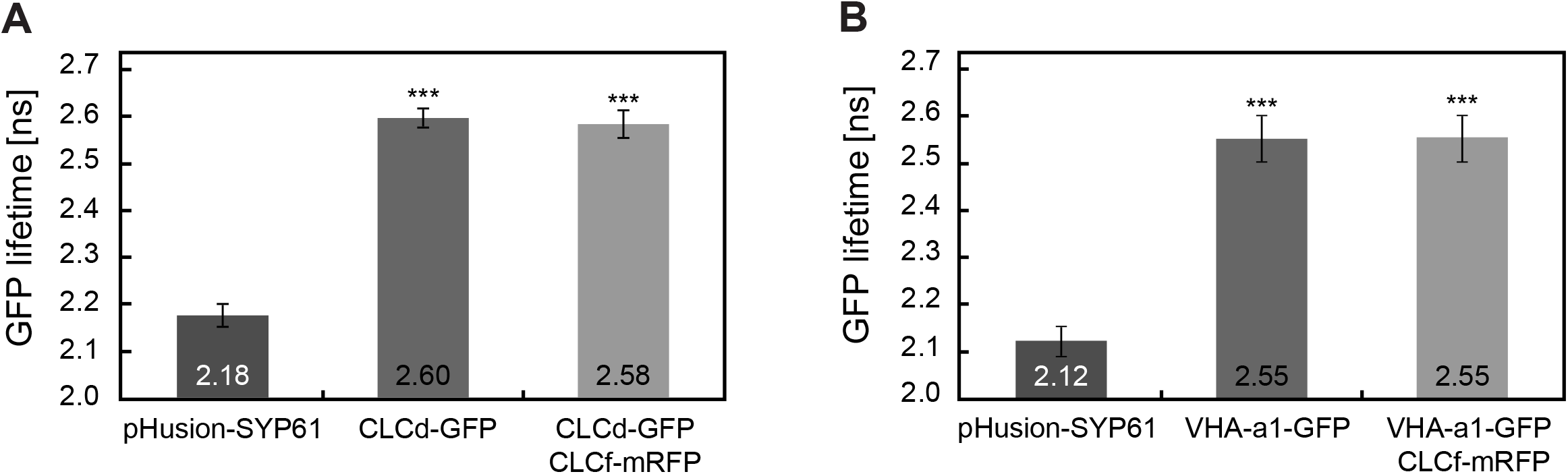
FRET-FLIM analysis. FRET-FLIM analyses was performed in cells of the root transition zone of 6-day-old Arabidopsis seedlings. **(A)** GFP fluorescence lifetime measurements in Arabidopsis lines stably expressing P16:pHusion-SYP61, UBQ10:CLCd-GFP or coexpressing UBQ10:CLCd-GFP and UBQ10:CLCf-mRFP. **(B)** GFP fluorescence lifetime measurements in Arabidopsis lines stably expressing P16:pHusion-SYP61, UBQ10:CLCd-GFP or coexpressing UBQ10:CLCd-GFP and UBQ10:CLCf-mRFP. Error bars indicate SD of 3 independent experiments. For each independent experiment 15 seedlings have been measured. Significant differences compared to positive the control SYP61-pHusion are indicated (*** indicate *p*-value < 0.001, Student’s t-test).

**Supplementary Figure S2:**
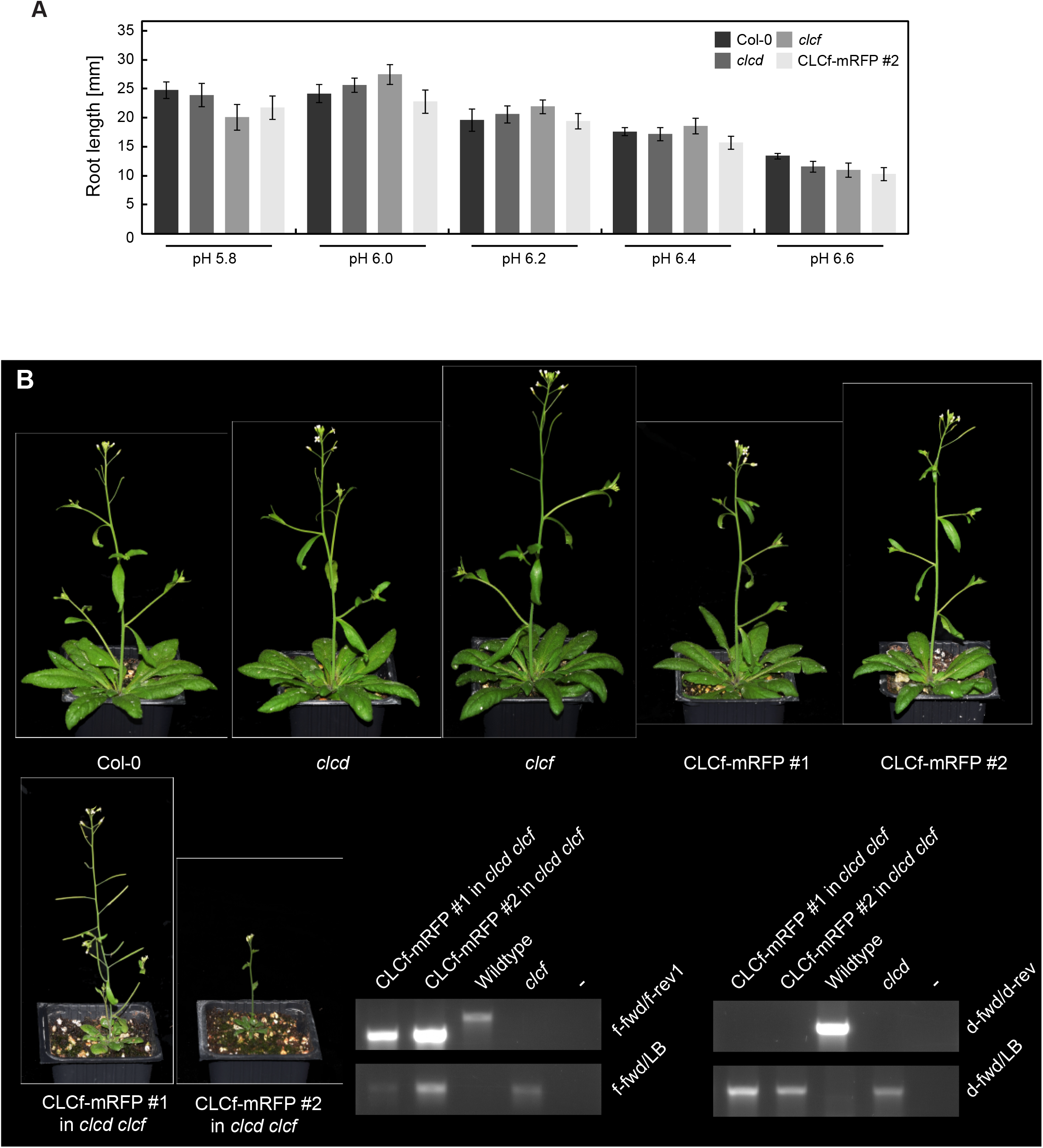
Phenotype of *clcd* and *clcf* single mutants and complementation of *clcd clcf*. **(A)** Root length measurements of 7-day-old Arabidopsis seedlings grown on chloride- and nitrate-free minimal medium with different pH values. Error bars indicate SE with n of 16 roots. **(B)** Phenotype of Arabidopsis wildtype Col-0, *clcd* and *clcf* single mutants, CLCf-mRFP #1, CLCf-mRFP #2, CLCf-mRFP #1 in *clcd clcf* and CLCf-mRFP #2 in *clcd clcf*. Plants have been grown in long-day conditions for 30. The presence of the *CLCf-mRFP* transgene and the respective T-DNA insertions in *clcf* and *clcd* were confirmed by genotyping.

**Supplementary Figure S3:**
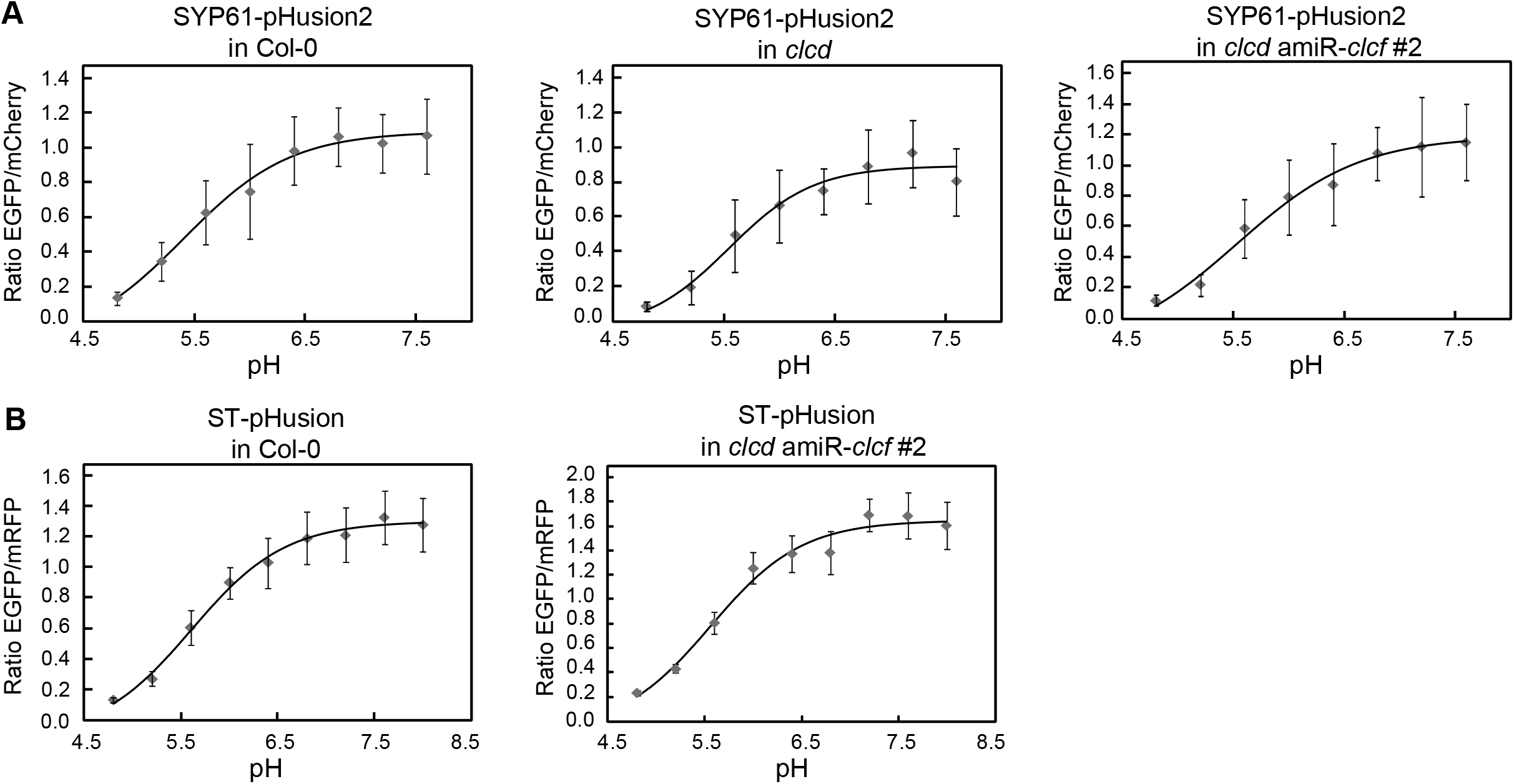
In vivo calibration curves. **(A)** In vivo calibration of UBQ10:SYP61-pHusion2 in wildtype Col-0, *clcd* and *clcd amiR-clcf #2*. **(B)** In vivo calibration of UBQ10:ST-pHusion in wild-type Col-0, *clcd* and *clcd amiR-clcf #2*. **(A-B)** Calibrations were performed in cells of the root elongation zone of 6-7-day-old Arabidopsis seedlings expressing the respective pH sensor. Error bars represent SD of n 15 seedlings per measurement.

## References

Accardi A, Miller C. 2004. Secondary active transport mediated by a prokaryotic homologue of ClC Cl-channels. Nature 427:803–807.

Anderson CT. 2016. We be jammin’: an update on pectin biosynthesis, trafficking and dynamics. Journal of Experimental Botany. doi:10.1093/jxb/erv501

Bassil E, Ohto M-A, Esumi T, Tajima H, Zhu Z, Cagnac O, Belmonte M, Peleg Z, Yamaguchi T, Blumwald E. 2011. The Arabidopsis intracellular Na+/H+ antiporters NHX5 and NHX6 are endosome associated and necessary for plant growth and development. Plant Cell 23:224–239.

Benková E, Michniewicz M, Sauer M, Teichmann T, Seifertová D, Jürgens G, Friml J. 2003. Local, efflux-dependent auxin gradients as a common module for plant organ formation. Cell 115:591– 602.

Bolte S, Cordelières FP. 2006. A guided tour into subcellular colocalization analysis in light microscopy. J Microsc 224:213–232.

Braun NA, Morgan B, Dick TP, Schwappach B. 2010. The yeast CLC protein counteracts vesicular acidification during iron starvation. J Cell Sci 123:2342–2350.

Brüx A, Liu T-Y, Krebs M, Stierhof Y-D, Lohmann JU, Miersch O, Wasternack C, Schumacher K. 2008. Reduced V-ATPase activity in the trans-Golgi network causes oxylipin-dependent hypocotyl growth Inhibition in Arabidopsis. Plant Cell 20:1088–1100.

Craft J, Samalova M, Baroux C, Townley H, Martinez A, Jepson I, Tsiantis M, Moore I. 2005. New pOp/LhG4 vectors for stringent glucocorticoid-dependent transgene expression in Arabidopsis. Plant J 41:899–918.

De Angeli A, Monachello D, Ephritikhine G, Frachisse JM, Thomine S, Gambale F, Barbier-Brygoo H. 2006. The nitrate/proton antiporter AtCLCa mediates nitrate accumulation in plant vacuoles. Nature 442:939–942.

Demes E, Besse L, Cubero-Font P, Satiat-Jeunemaitre B, Thomine S, De Angeli A. 2020. Dynamic measurement of cytosolic pH and [NO3-] uncovers the role of the vacuolar transporter AtCLCa in cytosolic pH homeostasis. Proc Natl Acad Sci U S A 117:15343–15353.

Dettmer J, Hong-Hermesdorf A, Stierhof Y-D, Schumacher K. 2006. Vacuolar H+-ATPase activity is required for endocytic and secretory trafficking in Arabidopsis. Plant Cell 18:715–730.

Dettmer J, Schubert D, Calvo-Weimar O, Stierhof Y-D, Schmidt R, Schumacher K. 2005. Essential role of the V-ATPase in male gametophyte development. Plant J 41:117–124.

Dragwidge JM, Scholl S, Schumacher K, Gendall AR. 2019. NHX-type Na+(K+)/H+ antiporters are required for TGN/EE trafficking and endosomal ion homeostasis in Arabidopsis thaliana. J Cell Sci 132. doi:10.1242/jcs.226472

French AP, Mills S, Swarup R, Bennett MJ, Pridmore TP. 2008. Colocalization of fluorescent markers in confocal microscope images of plant cells. Nat Protoc 3:619–628.

Grefen C, Donald N, Hashimoto K, Kudla J, Schumacher K, Blatt MR. 2010. A ubiquitin-10 promoter-based vector set for fluorescent protein tagging facilitates temporal stability and native protein distribution in transient and stable expression studies. Plant J 64:355–365.

Guo W, Zuo Z, Cheng X, Sun J, Li H, Li L, Qiu J-L. 2014. The chloride channel family gene CLCd negatively regulates pathogen-associated molecular pattern (PAMP)-triggered immunity in Arabidopsis. J Exp Bot 65:1205–1215.

Hajdukiewicz P, Svab Z, Maliga P. 1994. The small, versatile pPZP family of Agrobacterium binary vectors for plant transformation. Plant Mol Biol 25:989–994.

Hillmer S, Viotti C, Robinson DG. 2012. An improved procedure for low-temperature embedding of high-pressure frozen and freeze-substituted plant tissues resulting in excellent structural preservation and contrast. J Microsc 247:43–47.

Jentsch TJ, Pusch M. 2018. CLC Chloride Channels and Transporters: Structure, Function, Physiology, and Disease. Physiol Rev 98:1493–1590.

Jossier M, Kroniewicz L, Dalmas F, Le Thiec D, Ephritikhine G, Thomine S, Barbier-Brygoo H, Vavasseur A, Filleur S, Leonhardt N. 2010. The Arabidopsis vacuolar anion transporter, AtCLCc, is involved in the regulation of stomatal movements and contributes to salt tolerance. Plant J 64:563–576.

Kane PM, Jaskolka MC, Tuli F, Mitra C, Banerjee S, Tarsio M. 2020. Deciphering the isoform code contained in V o a-subunit isoforms of the V-ATPase. FASEB J 34:1–1.

Kasper D, Planells-Cases R, Fuhrmann JC, Scheel O, Zeitz O, Ruether K, Schmitt A, Poët M, Steinfeld R, Schweizer M, Kornak U, Jentsch TJ. 2005. Loss of the chloride channel ClC-7 leads to lysosomal storage disease and neurodegeneration. EMBO J 24:1079–1091.

Keinath NF, Kierszniowska S, Lorek J, Bourdais G, Kessler SA, Shimosato-Asano H, Grossniklaus U, Schulze WX, Robatzek S, Panstruga R. 2010. PAMP (Pathogen-associated Molecular Pattern)-induced Changes in Plasma Membrane Compartmentalization Reveal Novel Components of Plant Immunity *. J Biol Chem 285:39140–39149.

Krebs M, Held K, Binder A, Hashimoto K, Den Herder G, Parniske M, Kudla J, Schumacher K. 2012. FRET-based genetically encoded sensors allow high-resolution live cell imaging of Ca^+^ dynamics. Plant J 69:181–192.

Lampropoulos A, Sutikovic Z, Wenzl C, Maegele I, Lohmann JU, Forner J. 2013. GreenGate---a novel, versatile, and efficient cloning system for plant transgenesis. PLoS One 8:e83043.

Luo Y, Scholl S, Doering A, Zhang Y, Irani NG, Rubbo SD, Neumetzler L, Krishnamoorthy P, Van Houtte I, Mylle E, Bischoff V, Vernhettes S, Winne J, Friml J, Stierhof Y-D, Schumacher K, Persson S, Russinova E. 2015. V-ATPase activity in the TGN/EE is required for exocytosis and recycling in Arabidopsis. Nat Plants 1:15094.

Lupanga U, Röhrich R, Askani J, Hilmer S, Kiefer C, Krebs M, Kanazawa T, Ueda T, Schumacher K. 2020. The Arabidopsis V-ATPase is localized to the TGN/EE via a seed plant-specific motif. Elife 9. doi:10.7554/eLife.60568

Marmagne A, Vinauger-Douard M, Monachello D, de Longevialle AF, Charon C, Allot M, Rappaport F, Wollman F-A, Barbier-Brygoo H, Ephritikhine G. 2007. Two members of the Arabidopsis CLC (chloride channel) family, AtCLCe and AtCLCf, are associated with thylakoid and Golgi membranes, respectively. J Exp Bot 58:3385–3393.

Martinière A, Bassil E, Jublanc E, Alcon C, Reguera M, Sentenac H, Blumwald E, Paris N. 2013. In vivo intracellular pH measurements in tobacco and Arabidopsis reveal an unexpected pH gradient in the endomembrane system. Plant Cell 25:4028–4043.

McKay DW, Qu Y, McFarlane HE, Situmorang A, Gilliham M, Wege S. 2020. Cation Chloride Cotransporter 1 (CCC1) regulates pH and ionic conditions in the TGN/EE and is required for endomembrane trafficking. bioRxiv. doi:10.1101/2020.01.02.893073

Mindell JA. 2012. Lysosomal acidification mechanisms. Annu Rev Physiol 74:69–86.

Monachello D, Allot M, Oliva S, Krapp A, Daniel-Vedele F, Barbier-Brygoo H, Ephritikhine G. 2009. Two anion transporters AtClCa and AtClCe fulfil interconnecting but not redundant roles in nitrate assimilation pathways. New Phytol 183:88–94.

Nguyen CT, Agorio A, Jossier M, Depré S, Thomine S, Filleur S. 2016. Characterization of the Chloride Channel-Like, AtCLCg, Involved in Chloride Tolerance in Arabidopsis thaliana. Plant Cell Physiol 57:764–775.

Novarino G, Weinert S, Rickheit G, Jentsch TJ. 2010. Endosomal chloride-proton exchange rather than chloride conductance is crucial for renal endocytosis. Science 328:1398–1401.

Osei-Owusu J, Yang J, Leung KH, Ruan Z, Lü W, Krishnan Y, Qiu Z. 2021. Proton-activated chloride channel PAC regulates endosomal acidification and transferrin receptor-mediated endocytosis. Cell Rep 34:108683.

Pettersen EF, Goddard TD, Huang CC, Meng EC, Couch GS, Croll TI, Morris JH, Ferrin TE. 2021. UCSF ChimeraX: Structure visualization for researchers, educators, and developers. Protein Sci 30:70–82.

Raven JA. 2017. Chloride: essential micronutrient and multifunctional beneficial ion. J Exp Bot 38:359– 367.

Scheuring D, Viotti C, Krüger F, Künzl F, Sturm S, Bubeck J, Hillmer S, Frigerio L, Robinson DG, Pimpl P, Schumacher K. 2011. Multivesicular bodies mature from the trans-Golgi network/early endosome in Arabidopsis. Plant Cell 23:3463–3481.

Schindelin J, Arganda-Carreras I, Frise E, Kaynig V, Longair M, Pietzsch T, Preibisch S, Rueden C, Saalfeld S, Schmid B, Tinevez J-Y, White DJ, Hartenstein V, Eliceiri K, Tomancak P, Cardona A. 2012. Fiji: an open-source platform for biological-image analysis. Nat Methods 9:676–682.

Schrecker M, Korobenko J, Hite RK. 2020. Cryo-EM structure of the lysosomal chloride-proton exchanger CLC-7 in complex with OSTM1. Elife 9. doi:10.7554/eLife.59555

Schumacher K. 2014. pH in the plant endomembrane system-an import and export business. Curr Opin Plant Biol 22:71–76.

Schumacher K, Vafeados D, McCarthy M, Sze H, Wilkins T, Chory J. 1999. The Arabidopsis det3 mutant reveals a central role for the vacuolar H(+)-ATPase in plant growth and development. Genes Dev 13:3259–3270.

Schwab R, Ossowski S, Riester M, Warthmann N, Weigel D. 2006. Highly specific gene silencing by artificial microRNAs in Arabidopsis. Plant Cell 18:1121–1133.

Schwappach B, Stobrawa S, Hechenberger M, Steinmeyer K, Jentsch TJ. 1998. Golgi localization and functionally important domains in the NH2 and COOH terminus of the yeast CLC putative chloride channel Gef1p. J Biol Chem 273:15110–15118.

Shaner NC, Steinbach PA, Tsien RY. 2005. A guide to choosing fluorescent proteins. Nat Methods 2:905–909.

Stauber T, Jentsch TJ. 2013. Chloride in vesicular trafficking and function. Annu Rev Physiol 75:453– 477.

Sze H, Chanroj S. 2018. Plant Endomembrane Dynamics: Studies of K+/H+ Antiporters Provide Insights on the Effects of pH and Ion Homeostasis. Plant Physiol 177:875–895.

Teardo E, Frare E, Segalla A, De Marco V, Giacometti GM, Szabò I. 2005. Localization of a putative ClC chloride channel in spinach chloroplasts. FEBS Lett 579:4991–4996.

Ullrich F, Blin S, Lazarow K, Daubitz T, von Kries JP, Jentsch TJ. 2019. Identification of TMEM206 proteins as pore of PAORAC/ASOR acid-sensitive chloride channels. Elife 8. doi:10.7554/eLife.49187

Vasanthakumar T, Rubinstein JL. 2020. Structure and Roles of V-type ATPases. Trends Biochem Sci. doi:10.1016/j.tibs.2019.12.007

Viotti C, Bubeck J, Stierhof Y-D, Krebs M, Langhans M, van den Berg W, van Dongen W, Richter S, Geldner N, Takano J, Jürgens G, de Vries SC, Robinson DG, Schumacher K. 2010. Endocytic and secretory traffic in Arabidopsis merge in the trans-Golgi network/early endosome, an independent and highly dynamic organelle. Plant Cell 22:1344–1357.

von der Fecht-Bartenbach J, Bogner M, Dynowski M, Ludewig U. 2010. CLC-b-mediated NO-3/H+ exchange across the tonoplast of Arabidopsis vacuoles. Plant Cell Physiol 51:960–968.

von der Fecht-Bartenbach J, Bogner M, Krebs M, Stierhof Y-D, Schumacher K, Ludewig U. 2007. Function of the anion transporter AtCLC-d in the trans-Golgi network. Plant J 50:466–474.

Waadt R, Krebs M, Kudla J, Schumacher K. 2017. Multiparameter imaging of calcium and abscisic acid and high-resolution quantitative calcium measurements using R-GECO1-mTurquoise in Arabidopsis. New Phytol 216:303–320.

Wege S, Gilliham M, Henderson SW. 2017. Chloride: not simply a “cheap osmoticum”, but a beneficial plant macronutrient. J Exp Bot 68:3057–3069.

Wege S, Jossier M, Filleur S, Thomine S, Barbier-Brygoo H, Gambale F, De Angeli A. 2010. The proline 160 in the selectivity filter of the Arabidopsis NO(3)(-)/H(+) exchanger AtCLCa is essential for nitrate accumulation in planta. Plant J 63:861–869.

Weinert S, Gimber N, Deuschel D, Stuhlmann T, Puchkov D, Farsi Z, Ludwig CF, Novarino G, López-Cayuqueo KI, Planells-Cases R, Jentsch TJ. 2020. Uncoupling endosomal CLC chloride/proton exchange causes severe neurodegeneration. EMBO J 39:e103358.

Yang J, Zhang Y. 2015. I-TASSER server: new development for protein structure and function predictions. Nucleic Acids Res 43:W174–81.

Zhang GF, Driouich A, Staehelin LA. 1993. Effect of monensin on plant Golgi: re-examination of the monensin-induced changes in cisternal architecture and functional activities of the Golgi apparatus of sycamore suspension-cultured cells. J Cell Sci 104 (Pt 3):819–831.

Zhang Y, Nikolovski N, Sorieul M, Vellosillo T, McFarlane HE, Dupree R, Kesten C, Schneider R, Driemeier C, Lathe R, Lampugnani E, Yu X, Ivakov A, Doblin MS, Mortimer JC, Brown SP, Persson S, Dupree P. 2016. Golgi-localized STELLO proteins regulate the assembly and trafficking of cellulose synthase complexes in Arabidopsis. Nat Commun 7:11656.

Zheng H, Camacho L, Wee E, Batoko H, Legen J, Leaver CJ, Malhó R, Hussey PJ, Moore I. 2005. A Rab-E GTPase mutant acts downstream of the Rab-D subclass in biosynthetic membrane traffic to the plasma membrane in tobacco leaf epidermis. Plant Cell 17:2020–2036.

Zifarelli G, Pusch M. 2010. CLC transport proteins in plants. FEBS Lett 584:2122–2127.

